# Physiologically based pharmacokinetic (PBPK) modeling of the role of CYP2D6 polymorphism for metabolic phenotyping with dextromethorphan

**DOI:** 10.1101/2022.08.23.504981

**Authors:** Jan Grzegorzewski, Janosch Brandhorst, Matthias König

## Abstract

The cytochrome P450 2D6 (CYP2D6) is a key xenobiotic-metabolizing enzyme involved in the clearance of many drugs. Genetic polymorphisms in CYP2D6 contribute to the large inter-individual variability in drug metabolism and could affect metabolic phenotyping of CYP2D6 probe substances such as dextromethorphan (DXM). To study this question, we (i) established an extensive pharmacokinetics dataset for DXM; and (ii) developed and validated a physiologically based pharmacokinetic (PBPK) model of DXM and its metabolites dextrorphan (DXO) and dextrorphan O-glucuronide (DXO-Glu) based on the data. Drug-gene interactions (DGI) were introduced by accounting for changes in CYP2D6 enzyme kinetics depending on activity score (AS), which in combination with AS for individual polymorphisms allowed us to model CYP2D6 gene variants. Variability in CYP3A4 and CYP2D6 activity was modeled based on in vitro data from human liver microsomes. Model predictions are in very good agreement with pharmacokinetics data for CYP2D6 polymorphisms, CYP2D6 activity as described by the AS system, and CYP2D6 metabolic phenotypes (UM, EM, IM, PM). The model was applied to investigate the genotype-phenotype association and the role of CYP2D6 polymorphisms for metabolic phenotyping using the urinary cumulative metabolic ratio (UCMR), DXM/(DXO+DXO-Glu). The effect of parameters on UCMR was studied via sensitivity analysis. Model predictions indicate very good robustness against the intervention protocol (i.e. application form, dosing amount, dissolution rate, and sampling time) and good robustness against physiological variation. The model is capable of estimating the UCMR dispersion within and across populations depending on activity scores. Moreover, the distribution of UCMR and the risk of genotype-phenotype mismatch could be estimated for populations with known CYP2D6 genotype frequencies. The model can be applied for individual prediction of UCMR and metabolic phenotype based on CYP2D6 genotype. Both, model and database are freely available for reuse.

## 1 INTRODUCTION

The cytochrome P450 (CYPs) superfamily of enzymes has a central role in the clearance of many substances and drugs, with the isoform 2D6 (CYP2D6) being one of the most important xenobiotic-metabolizing enzymes. CYP2D6 is involved in the clearance of around 20% of the most prescribed drugs (Saravanakumar et al., 2019) including antiarrhythmics having a small therapeutic range (e.g. flecainide, procainamide, mexiletine), anticancer agents (e.g. tamoxifen), antidepressants (e.g. citalopram, fluoxetine, duloxetine: venlafaxine), antipsychotics (e.g. aripiprazole, haloperidol, thioridazine), *β*-blockers (metoprolol), analgesics (tramadol, oxycodone, codeine), and antitussives (dextromethorphan) (Kibaly et al., 2021; Hurtado et al., 2020). CYP2D6-mediated drug response exhibits a particularly large inter-individual variability which poses a challenge for personalized dosage of medication by underdosing on the one hand and toxic side effects on the other. The activity of CYP2D6 is known to be majorly dependent on genetic variants (Preskorn et al., 2013; Berm et al., 2013; Shah and Smith, 2015) with polymorphism of CYP2D6 being related to the risk of adverse effects, non-response during treatment, and death by drug intoxication (Gasche et al., 2004; Kawanishi et al., 2004; Rau et al., 2004; Zackrisson et al., 2010).

In the late 70s, a polymorphism in debrisoquine hydroxylation (Mahgoub et al., 1977) and sparteine oxidation (Eichelbaum et al., 1979) was discovered and subsequently attributed to allelic variants of the CYP2D6 gene. In the following years, CYP2D6 became one of the most studied drug-metabolizing enzymes. Genetic variants were classified into distinct phenotypes and subjects carrying combinations of these variants were categorized as poor metabolizer (gPM), intermediate metabolizer (gIM), extensive metabolizer (gEM), and ultra rapid metabolizer (gUM) (Zanger et al., 2004; Gaedigk et al., 2017). This classification is based on the relationship between genetic variants and CYP2D6-mediated drug response. For these genetically predicted phenotypes, we use the ‘g’ nomenclature as they can be easily confused with the actual *in vivo* metabolic phenotype, determined based on pharmacokinetic measurements after the administration of CYP2D6 test drugs. Nowadays, the CYP2D6 activity score (AS) system, a more refined metric, is often applied to characterize genotype-phenotype associations (Gaedigk et al., 2018a). In the system, discrete values between 0 and 1 are assigned to gene variants. The final activity score is calculated by the sum of the activity scores of both alleles. For instance, a person with diplotype *1/*3 (the variant *1 has an AS of 1 and the variant *3 has no activity with an AS of 0) has an overall AS of 1. Higher activity scores than 2 and additional complexity arise from copy number variation (CNV), chimeras, and hybrids with the pseudo gene CYP2D7. This can result in ambiguities and difficulties in the assignment of the correct diplotype and activity score (Gaedigk et al., 2007; Nofziger and Paulmichl, 2018; Gaedigk et al., 2019). Of note, AS specifics are still under heavy debate and regularly updated (Caudle et al., 2020). A multitude of population studies have been conducted to identify and associate allele variants with metabolic phenotypes within and across populations (Gaedigk et al., 2017). Over 130 CYP2D6 star (*) allele haplotypes have been identified and subsequently cataloged by the Pharmacogene Variation (PharmVar) Consortium into PharmGKB with their respective activity score contribution (Gaedigk et al., 2018b; Whirl-Carrillo et al., 2021).

Various methods exist for the metabolic phenotyping based on test substances. The gold standard is plasma concentration sampling of probe substances and their metabolites at various time points after the administration. (Partial) clearance rates and the relative enzyme activities can be calculated from these plasma time profiles. Simplified methods have been established for many probe substances which do not require repeated sampling of blood, e.g., the (cumulative) metabolic ratios between the probe substance and one or several of its metabolites at a single time point in blood, plasma, or urine are utilized as such proxy measures. Large-scale population studies often tend to employ urinary ratios of metabolites. Alternatively, sampling of saliva and breath are worth considering (De Kesel et al., 2016). Probe substances for metabolic phenotyping of CYP2D6 are debrisoquine, dextromethorphan, metoprolol, or sparteine (Frank et al., 2007; Fuhr et al., 2007). Bufuralol is less popular but well suited for *in vitro* investigations due to its fluorescent properties (Zanger et al., 2004). Although debrisoquine and sparteine have excellent properties for CYP2D6 phenotyping, they have been withdrawn from clinical use in most countries and are therefore no longer readily available. Frequently in use for the phenotyping of CYP2D6 activity are metoprolol and dextromethorphan.

Dextromethorphan (DXM) is an over-the-counter, antitussive, non-narcotic, synthetic analog of codeine affecting the activity of numerous channels and receptors in the brain that trigger the cough reflex (Silva and Dinis-Oliveira, 2020). It is generally well-tolerated, considered safe in therapeutic dosage, and easily available (Fuhr et al., 2007). Besides therapeutic purposes, DXM is most commonly applied as a probe substance for CY2D6 phenotyping, alone or with other probe substances in a cocktail. DXM can be administered orally and intravenously, has low bioavailability (≈50%) and a rapid first-pass effect in the intestine and liver. Typically only about half of the dose is recovered in urine over at least 12 hours after administration, primarily as glucuronides (Schadel et al., 1995; Capon et al., 1996; Tennezé et al., 1999; Strauch et al., 2009). In the systemic circulation, ≈55-65% of DXM is non-specifically bound to plasma proteins (Lutz and Isoherranen, 2012; Taylor et al., 2016).

The biotransformation of DXM is mostly confined to the liver, where DXM is O-demethylated by CYP2D6 to the active metabolite dextrorphan (DXO). Subsequently to O-demethylation, most of the DXO is rapidly transformed via UDP-glucuronosyltransferase (UGT) to dextrorphan O-glucuronide (DXO-Glu) and excreted via the urine. In individuals without any functional variant of CYP2D6, the metabolization of DXM to DXO is extremely slow but still present. Apparently, the O-demethylation is not exclusively mediated by CYP2D6, and it has been demonstrated *in vitro* that O-demethylation of DXM can be marginally mediated by CYP3A4, CYP3A5 and CYP2C9 (von Moltke et al., 1998; McGinnity et al., 2000; Takashima et al., 2005; Yu and Haining, 2001). In line with this observation, inhibition of CYP2D6, e.g., barely affects poor metabolizer (Pope et al., 2004). The second pathway of DXM metabolization goes via N-demethylation to 3-methoxymorphinan which is mainly catalyzed via CYP3A4. Subsequently, 3-methoxymorphinan and DXO are biotransformed to 3-hydroxymorphinan which is then rapidly transformed via glucuronidation to hydroxymorphian O-glucuronide and excreted in the urine. The urinary cumulative metabolic ratio (UCMR) of DXM to its metabolites DXM/(DXO+DXO-Glu) is a widely applied measure for the *in vivo* CYP2D6 phenotyping.

An crucial question for metabolic phenotyping and liver function testing is how CYP2D6 polymorphisms affect the pharmacokinetics of DXM and metabolic phenotyping based on DXM, such as the UCMR. The objective of this work was to answer this question by the means of physiologically based pharmacokinetic (PBPK) modeling of DXM.

## 2 MATERIAL AND METHODS

### 2.1 Pharmacokinetics database of DXM

Pharmacokinetics data of DXM was systematically curated from literature for model development, parameterization, and validation. Curation efforts were mainly focused on concentration-time profiles of DXM, DXO, and DXO-Glu in plasma or serum and their amounts or ratios in urine. The data is accompanied by metadata on the investigated subjects and groups (e.g. CYP2D6 genotype or activity score) and the applied intervention (e.g. dose and application form of DXM). All data was curated using an established curation pipeline (Grzegorzewski et al., 2022) and is available via the pharmacokinetics database PK-DB (https://pk-db.com) (Grzegorzewski et al., 2021). As a first step, a PubMed search for the pharmacokinetics of dextromethorphan in combination with genotyping and/or phenotyping was performed with the search query https://pubmed.ncbi.nlm.nih.gov/?term=dextromethorphan+AND+%28phenotype+OR+phenotyping%29+AND+genotype. The literature corpus was extended with drug cocktail studies from PK-DB (Grzegorzewski et al., 2022), secondary literature from references, and results from PKPDAI with the search query https://app.pkpdai.com/?term=dextromethorphan Gonzalez Hernandez et al. (2021). Data was selected and curated based on eligibility criteria, see below. During the curation process, the initial corpus was updated by additional publications from the references and citations. A subset of the studies only reported pharmacokinetic parameters without timecourses. These studies were curated but not further used in the following analyses.

To be eligible, studies had to report in vivo pharmacokinetics data for adult (age ≥ 18) humans after administration of DXM or DXM hydrobromide. The application route of DXM was restricted to oral (PO) or intravenous (IV). All application forms (e.g. tablet, capsule, solution) were accepted. No restrictions were imposed on the dosing amount of DXM or coadministrations of other substances. Studies containing coadminstrations that inhibit or induce the pharmacokinetics of DXM were identified during the modeling process and excluded. The relevant outcome measures are concentration-time profiles in plasma, serum, and urine amounts of DXM, DXM metabolites, or metabolic ratios of metabolites such as UCMR. Studies containing pharmacokinetic parameters of DXM and its metabolites (e.g. clearance, half-life, AUC) and (urinary cumulative) metabolic ratios of DXM and its metabolites were included. Data containing timecourses and CYP2D6 genotype information were prioritized. Non-healthy subjects were excluded if the disease is known to affect the pharmacokinetics of DXM or DXM metabolites. Study B from the PhD thesis of Frank (2009) highly deviates from the remaining data and was therefore excluded. Further, Wyen et al. (2008) was identified as a duplicate of Study E from the PhD thesis of Frank (2009) and excluded. The final set of curated studies used in the presented analyses is provided in Tab. 1.

**Table 1.**
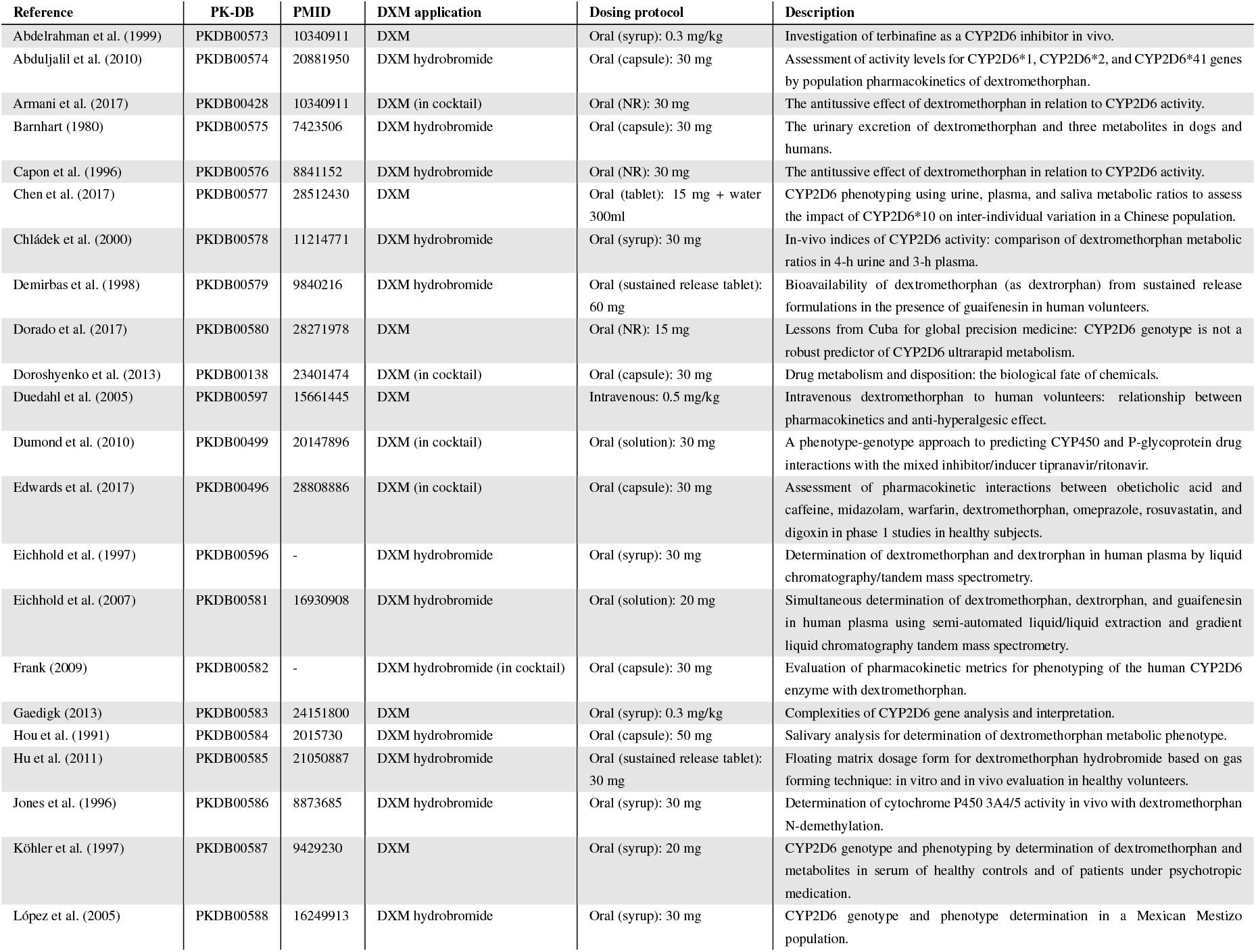
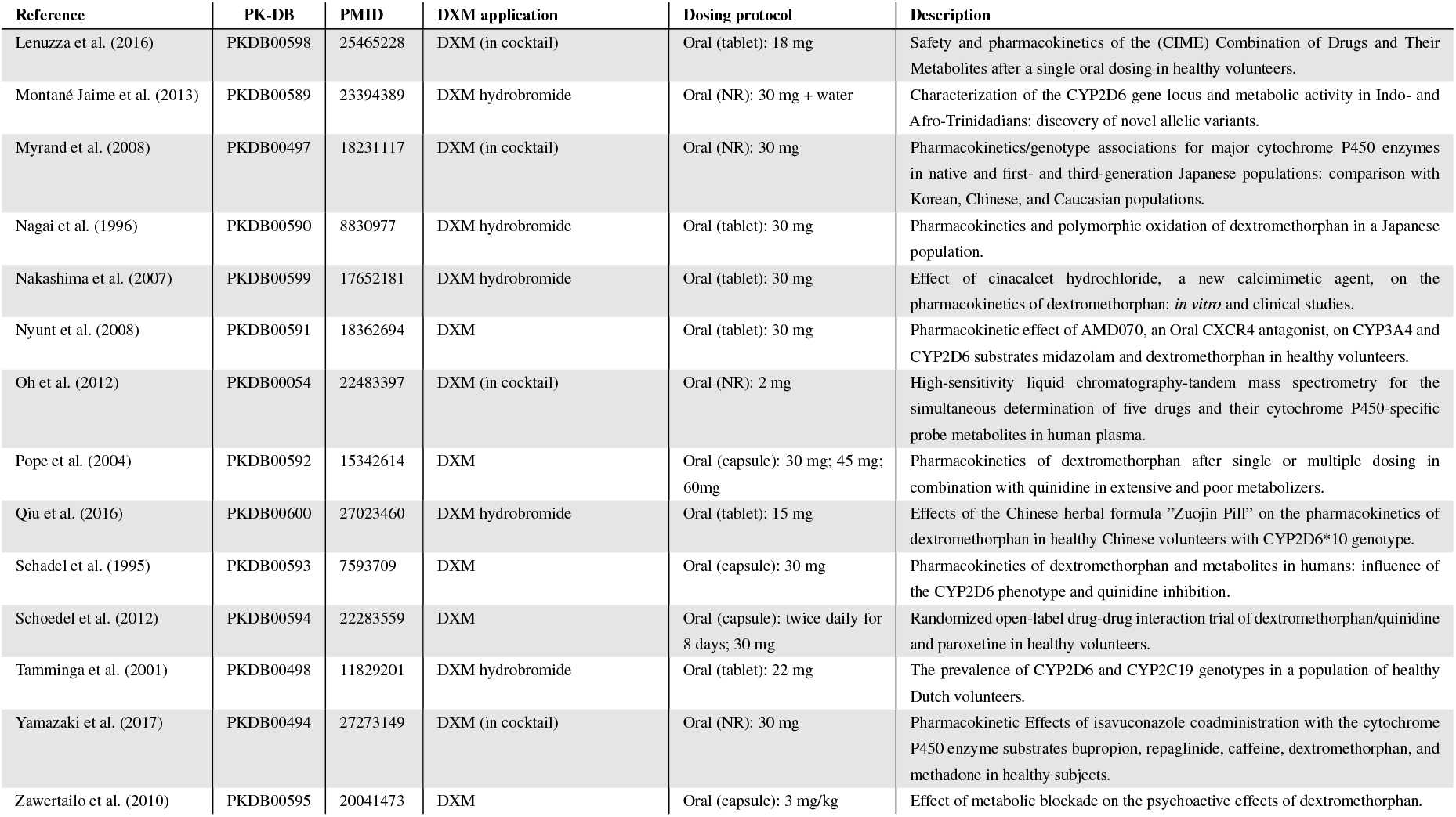
Clinical studies with pharmacokinetics used for model evaluation. NR: not reported, DXM: dextromethorphan.

| Reference | PK-DB | PMID | DXM application | Dosing protocol | Description |
| --- | --- | --- | --- | --- | --- |
| Abdelrahman et al. (1999) | PKDB00573 | 10340911 | DXM | Oral (syrup): 0.3 mg/kg | Investigation of terbinafine as a CYP2D6 inhibitor in vivo. |
| Abduljalil et al. (2010) | PKDB00574 | 20881950 | DXM hydrobromide | Oral (capsule): 30 mg | Assessment of activity levels for CYP2D6*1, CYP2D6*2, and CYP2D6*41 genes by population pharmacokinetics of dextromethorphan. |
| Armani et al. (2017) | PKDB00428 | 10340911 | DXM (in cocktail) | Oral (NR): 30 mg | The antitussive effect of dextromethorphan in relation to CYP2D6 activity. |
| Barnhart (1980) | PKDB00575 | 7423506 | DXM hydrobromide | Oral (capsule): 30 mg | The urinary excretion of dextromethorphan and three metabolites in dogs and humans. |
| Capon et al. (1996) | PKDB00576 | 8841152 | DXM hydrobromide | Oral (NR): 30 mg | The antitussive effect of dextromethorphan in relation to CYP2D6 activity. |
| Chen et al. (2017) | PKDB00577 | 28512430 | DXM | Oral (tablet): 15 mg + water 300ml | CYP2D6 phenotyping using urine, plasma, and saliva metabolic ratios to assess the impact of CYP2D6*10 on inter-individual variation in a Chinese population. |
| Chládek et al. (2000) | PKDB00578 | 11214771 | DXM hydrobromide | Oral (syrup): 30 mg | In-vivo indices of CYP2D6 activity: comparison of dextromethorphan metabolic ratios in 4-h urine and 3-h plasma. |
| Demirbas et al. (1998) | PKDB00579 | 9840216 | DXM hydrobromide | Oral (sustained release tablet): 60 mg | Bioavailability of dextromethorphan (as dextrorphan) from sustained release formulations in the presence of guaifenesin in human volunteers. |
| Dorado et al. (2017) | PKDB00580 | 28271978 | DXM | Oral (NR): 15 mg | Lessons from Cuba for global precision medicine: CYP2D6 genotype is not a robust predictor of CYP2D6 ultrarapid metabolism. |
| Doroshenko et al. (2013) | PKDB00138 | 23401474 | DXM (in cocktail) | Oral (capsule): 30 mg | Drug metabolism and disposition: the biological fate of chemicals. |
| Duedahl et al. (2005) | PKDB00597 | 15661445 | DXM | Intravenous: 0.5 mg/kg | Intravenous dextromethorphan to human volunteers: relationship between pharmacokinetics and anti-hyperalgesic effect. |
| Dumond et al. (2010) | PKDB00499 | 20147896 | DXM (in cocktail) | Oral (solution): 30 mg | A phenotype-genotype approach to predicting CYP450 and P-glycoprotein drug interactions with the mixed inhibitor/inducer tipranavir/ritonavir. |
| Edwards et al. (2017) | PKDB00496 | 28808886 | DXM (in cocktail) | Oral (capsule): 30 mg | Assessment of pharmacokinetic interactions between obeticholic acid and caffeine, midazolam, warfarin, dextromethorphan, omeprazole, rosuvastatin, and digoxin in phase I studies in healthy subjects. |
| Eichhold et al. (1997) | PKDB00596 | - | DXM hydrobromide | Oral (syrup): 30 mg | Determination of dextromethorphan and dextrorphan in human plasma by liquid chromatography/tandem mass spectrometry. |
| Eichhold et al. (2007) | PKDB00581 | 16930908 | DXM hydrobromide | Oral (solution): 20 mg | Simultaneous determination of dextromethorphan, dextrorphan, and guaifenesin in human plasma using semi-automated liquid/liquid extraction and gradient liquid chromatography tandem mass spectrometry. |
| Frank (2009) | PKDB00582 | - | DXM hydrobromide (in cocktail) | Oral (capsule): 30 mg | Evaluation of pharmacokinetic metrics for phenotyping of the human CYP2D6 enzyme with dextromethorphan. |
| Gaedigk (2013) | PKDB00583 | 24151800 | DXM | Oral (syrup): 0.3 mg/kg | Complexities of CYP2D6 gene analysis and interpretation. |
| Hou et al. (1991) | PKDB00584 | 2015730 | DXM hydrobromide | Oral (capsule): 50 mg | Salivary analysis for determination of dextromethorphan metabolic phenotype. |
| Hu et al. (2011) | PKDB00585 | 21050887 | DXM hydrobromide | Oral (sustained release tablet): 30 mg | Floating matrix dosage form for dextromethorphan hydrobromide based on gas forming technique: in vitro and in vivo evaluation in healthy volunteers. |
| Jones et al. (1996) | PKDB00586 | 8873685 | DXM hydrobromide | Oral (syrup): 30 mg | Determination of cytochrome P450 3A4/5 activity in vivo with dextromethorphan N-demethylation. |
| Köhler et al. (1997) | PKDB00587 | 9429230 | DXM | Oral (syrup): 20 mg | CYP2D6 genotype and phenotyping by determination of dextromethorphan and metabolites in serum of healthy controls and of patients under psychotropic medication. |
| López et al. (2005) | PKDB00588 | 16249913 | DXM hydrobromide | Oral (syrup): 30 mg | CYP2D6 genotype and phenotype determination in a Mexican Mestizo population. |
| Lenuzza et al. (2016) | PKDB00598 | 25465228 | DXM (in cocktail) | Oral (tablet): 18 mg | Safety and pharmacokinetics of the (CIME) Combination of Drugs and Their Metabolites after a single oral dosing in healthy volunteers. |
| Montané Jaime et al. (2013) | PKDB00589 | 23394389 | DXM hydrobromide | Oral (NR): 30 mg + water | Characterization of the CYP2D6 gene locus and metabolic activity in Indo- and Afro-Trinidadians: discovery of novel allelic variants. |
| Myrand et al. (2008) | PKDB00497 | 18231117 | DXM (in cocktail) | Oral (NR): 30 mg | Pharmacokinetics/genotype associations for major cytochrome P450 enzymes in native and first- and third-generation Japanese populations: comparison with Korean, Chinese, and Caucasian populations. |
| Nagai et al. (1996) | PKDB00590 | 8830977 | DXM hydrobromide | Oral (tablet): 30 mg | Pharmacokinetics and polymorphic oxidation of dextromethorphan in a Japanese population. |
| Nakashima et al. (2007) | PKDB00599 | 17652181 | DXM hydrobromide | Oral (tablet): 30 mg | Effect of cinacalcet hydrochloride, a new calcimimetic agent, on the pharmacokinetics of dextromethorphan: *in vitro* and clinical studies. |
| Nyunt et al. (2008) | PKDB00591 | 18362694 | DXM | Oral (tablet): 30 mg | Pharmacokinetic effect of AMD070, an Oral CXCR4 antagonist, on CYP3A4 and CYP2D6 substrates midazolam and dextromethorphan in healthy volunteers. |
| Oh et al. (2012) | PKDB00054 | 22483397 | DXM (in cocktail) | Oral (NR): 2 mg | High-sensitivity liquid chromatography-tandem mass spectrometry for the simultaneous determination of five drugs and their cytochrome P450-specific probe metabolites in human plasma. |
| Pope et al. (2004) | PKDB00592 | 15342614 | DXM | Oral (capsule): 30 mg; 45 mg; 60mg | Pharmacokinetics of dextromethorphan after single or multiple dosing in combination with quinidine in extensive and poor metabolizers. |
| Qiu et al. (2016) | PKDB00600 | 27023460 | DXM hydrobromide | Oral (tablet): 15 mg | Effects of the Chinese herbal formula "Zuojin Pill" on the pharmacokinetics of dextromethorphan in healthy Chinese volunteers with CYP2D6*10 genotype. |
| Schadel et al. (1995) | PKDB00593 | 7593709 | DXM | Oral (capsule): 30 mg | Pharmacokinetics of dextromethorphan and metabolites in humans: influence of the CYP2D6 phenotype and quinidine inhibition. |
| Schoedel et al. (2012) | PKDB00594 | 22283559 | DXM | Oral (capsule): twice daily for 8 days; 30 mg | Randomized open-label drug-drug interaction trial of dextromethorphan/quinidine and paroxetine in healthy volunteers. |
| Tamminga et al. (2001) | PKDB00498 | 11829201 | DXM hydrobromide | Oral (tablet): 22 mg | The prevalence of CYP2D6 and CYP2C19 genotypes in a population of healthy Dutch volunteers. |
| Yamazaki et al. (2017) | PKDB00494 | 27273149 | DXM (in cocktail) | Oral (NR): 30 mg | Pharmacokinetic Effects of isavuconazole coadministration with the cytochrome P450 enzyme substrates bupropion, repaglinide, caffeine, dextromethorphan, and methadone in healthy subjects. |
| Zawertailo et al. (2010) | PKDB00595 | 20041473 | DXM | Oral (capsule): 3 mg/kg | Effect of metabolic blockade on the psychoactive effects of dextromethorphan. |

For the selection and evaluation of studies from the literature, the PRISMA-ScR guidelines were adopted where applicable (Tricco et al., 2018). The initial corpus contained 404 studies. After screening, application of eligibility criteria, and prioritization, a total of 47 studies were curated (see supplementary Fig. S1). Of these studies, 36 contained data used in the present work (Tab. 1).

### 2.2 PBPK model of DXM

The PBPK model of DXM, DXO, and DXO-Glu (Fig. 1) was encoded in the Systems Biology Markup Language (SBML) (Hucka et al., 2019; Keating et al., 2020). For development and visualization, sbmlutils (König, 2021b) and cy3sbml (König et al., 2012; König and Rodriguez, 2019) were used. The model utilizes ordinary differential equations (ODE) which were numerically solved by sbmlsim (König, 2021a) based on the high-performance SBML simulator libroadrunner (Somogyi et al., 2015; Welsh et al., 2022). It is available in SBML under CC-BY 4.0 license from https://github.com/matthiaskoenig/dextromethorphan-model. Within this work, version 0.9.5 of the model was used Grzegorzewski and König (2022).

**Figure 1.**
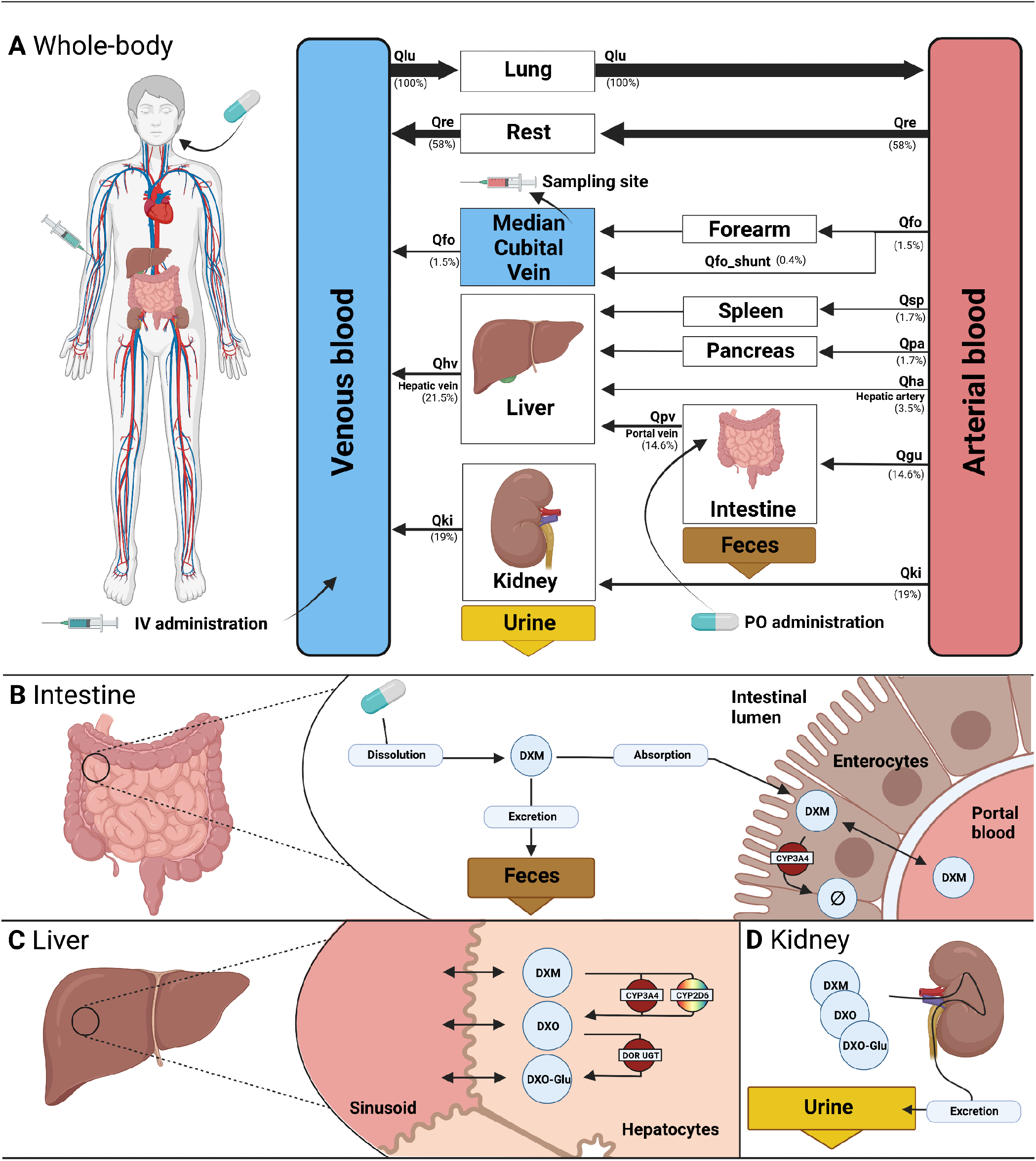
PBPK model of dextromethorphan (DXM). **A)** whole body model consisting of liver, kidney, intestine, forearm, lung and the rest compartment. Organs with minor relevance are not modeled explicitly and lumped into the rest compartment. Organs are coupled via the systemic circulation with arrow width proportional to relative blood flow. DXM can be administered intravenously (IV) or orally (PO) with DXM appearing in the venous blood or intestine, respectively. **B)** intestine model consisting of dissolution, absorption and excretion of DXM. Only a fraction of DXM is absorbed with the remainder excreted in the feces. First pass metabolism of DXM via CYP3A4 in the intestine reduces the amount of DXM appearing in the circulation. **C)** liver model consisting of DXM → DXO conversion via CYP2D6 and CYP3A4 and subsequent glucuronidation to DXO-Glu. **D** kidney model for the urinary excretion of DXM, DXO and DXO-Glu. Created with BioRender.com.

The model is hierarchically organized with submodels coupled using hierarchical model composition (Smith et al., 2015). The top layer represents the whole body with organs and tissues connected via the blood flow. The lower layer describes substance-related processes within the tissues. Tissues with minor influence on the pharmacokinetics of DXM, DXO, or DXO-Glu are lumped into the ‘rest’ compartment. Intravenous and oral application of DXM appears in the venous and intestinal compartments, respectively. A fraction of DXM is absorbed via the intestinal wall into the systemic circulation. The remainder is excreted via the feces. The plasma concentration is evaluated at the median cubital vein.

The distribution of DXM, DXO, and DXO-Glu between plasma and tissue compartments is based on tissue-to-plasma partition coefficients (K_p_) and the corresponding rates of tissue distribution (f_tissue_).

The metabolism of DXM only includes processes relevant for the simulation of the reported pharmacokinetics data (see Fig. 1B and C). Routes of minor contribution such as N-demethylation of DXM in the liver were neglected. Metabolic reactions take place in the intestine and liver and are modeled using irreversible Michaelis-Menten reaction kinetics of the form 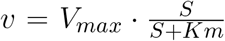, with V_max_ and K_m_ for CYP3A4 and CYP2D6 sampled from distributions as described below. The conversion of DXM to DXO can be either catalyzed via CYP2D6 (main route) or CYP3A4 (minor route) in the liver. Reactions with other products than DXM, DXO, and DXO-Glu were modeled as annihilation, i.e. the products of the reaction are not modeled explicitly. DXM, DXO, and DXO-Glu are eliminated into the urine via renal excretion.

A subset of model parameters was fitted by minimizing the distance between model predictions and subsets of the data in Fig. 4, Fig. 5, Fig. 6, Fig. 7, Fig. 8, and Fig. 9.

### 2.3 CYP3A4 and CYP2D6

CYP3A4 and CYP2D6 variability was modeled via correlated bivariate lognormal distributions fitted to in *vitro* data for CYP2D6 (Yang et al., 2012; Storelli et al., 2019a) and CYP3A4 (Yang et al., 2012), respectively. The data was log10 transformed and a Gaussian, parameterized by the mean (*μ*) and standard deviation, was fitted by maximum likelihood estimation. The multivariate distribution was realized by a Gaussian copula which in turn was parameterized by Kendall’s tau correlation coefficient from the data (see Fig. 2 for data and model).

**Figure 2.**
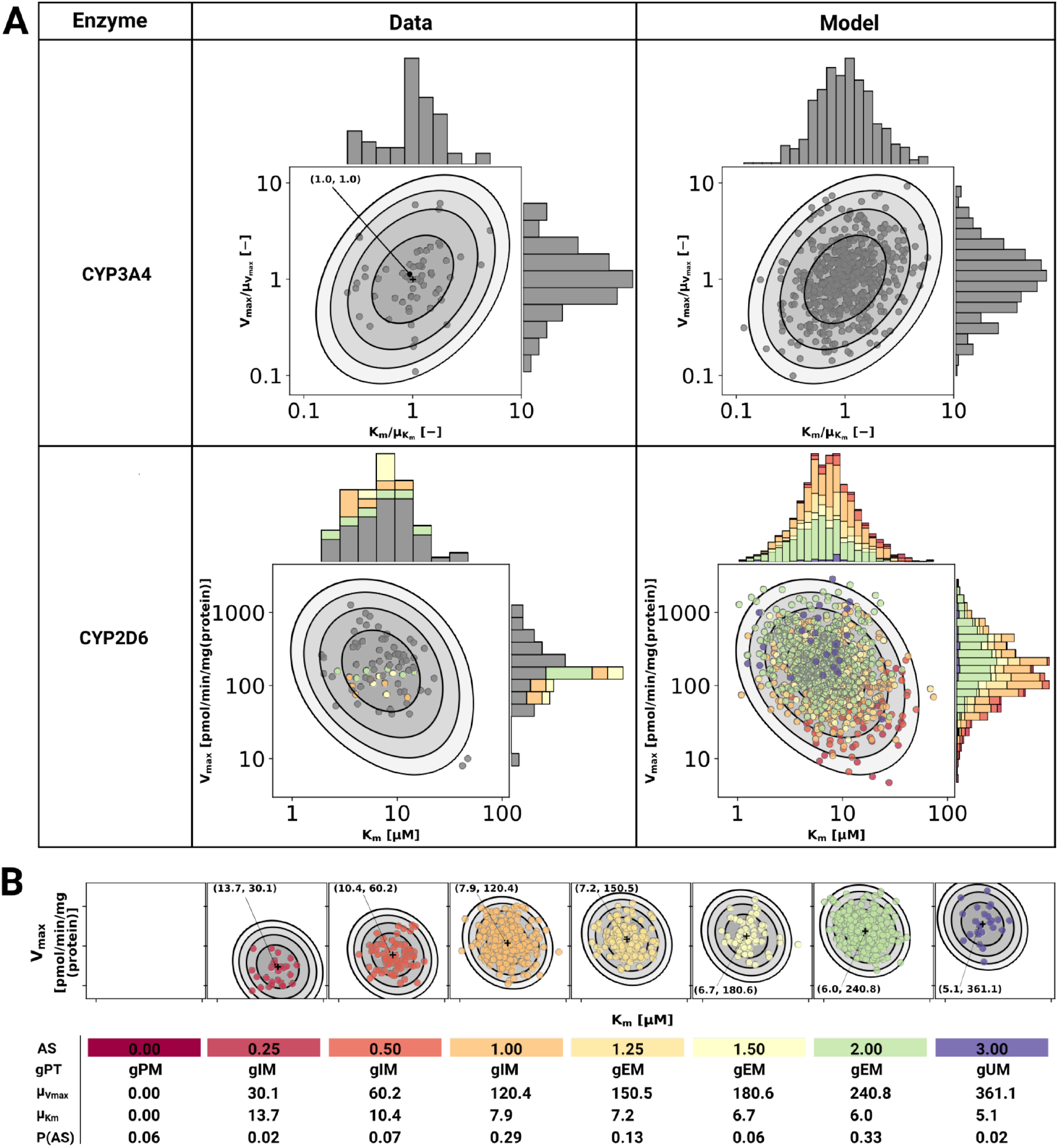
Model of CYP3A4 and CYP2D6. **A)** CYP3A4 and CYP2D6 distributions. Conversion of DXM → DXO via CYP3A4 and CYP2D6 are modeled via Michaelis-Menten kinetics. Variability was included via two-dimensional lognormal distributions of Michaelis-Menten coefficient (K_m_) and maximum rate of reaction (V_max_). The distribution parameters were determine by fitting to *in vitro* data in human liver microsomes. Variability of CYP3A4 parameters was measured by midazolam (Yang et al., 2012), variability of CYP2D6 parameters via DXM (Yang et al., 2012; Storelli et al., 2019a). To transfer the CYP3A4 data from midazolam to DXM normalized values were used. The distribution of CYP2D6 was modeled as a mixture model of the underlying activity scores as depicted in B. The model CYP3A4 and CYP2D6 distributions were sampled with each point corresponding to a combination of V_max_ and K_m_. CYP2D6 data was color-coded by the respective activity score. **B)** CYP2D6 activity score model. CYP2D6 activity was modeled via a mixture model of individual activity scores. With increasing activity score the V_max_ for the DXM → DXO conversion increases and the *μ*_Km_ for DXM decreases, i.e., reaction velocity and affinity for the substrate increase. The table provides AS, genetic phenotype (gPT), mean V_max_, mean K_m_, and AS frequency in curated UCMR data (P(AS)). In case of AS=0.0 CYP2D6 has no activity for the DXM → DXO conversion.

In order to model the effect of the CYP2D6 AS on the activity, V_max_ was assumed to be proportional to the AS, V_max_ α AS and K_m_ was scaled by the activity score along the first principle component of log10(K_m_) and log10(V_max_) (principal component regression). To model the effect of genetic polymorphisms of CYP2D6, pharmacogenetic variants in the CYP2D6 gene were mapped to their AS and the total activity calculated as the sum of the activity of the two alleles. The genotype-phenotype definitions (i.e. allele variant to AS mapping) were used from PharmGKB (https://www.pharmgkb.org/page/cyp2d6RefMaterials, accessed on 2022-01-10) (Whirl-Carrillo et al., 2021) (Tab. S1).

The stochastic model of CYP2D6 kinetics for a given population consists of a mixture model comprised from the models for each AS weighted by their respective frequency P(AS), i.e., 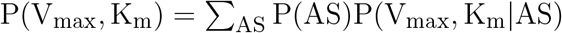. To simulate a given AS, the respective K_m_ and V_max_ values were used (see Fig. 2). The variability in pharmacokinetics was simulated by sampling K_m_ and V_max_ from the CYP3A4 and CYP2D6 distributions. Distributions of CYP3A4 and CYP2D6 parameters were assumed to be statistically independent. To simulate different populations, the AS frequencies for the respective biogeographical population were used from PharmGKB (https://www.pharmgkb.org/page/cyp2d6RefMaterials, accessed on 2022-01-10) (Whirl-Carrillo et al., 2021) (Tab. S2).

### 2.4 CYP2D6 metabolic phenotype

The metabolic phenotypes ultrarapid metabolizer (UM), extensive metabolizer (EM), intermediate metabolizer (IM), and poor metabolizer (PM) were assigned based on the urinary cumulative metabolic ratio of DXM to total dextrorphan 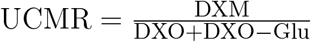 with the following cutoffs: PM: UCMR ≥ 0.3, IM: 0.03 ≤ UCMR < 0.3, EM: 0.0003 ≤ UCMR < 0.03, UM: UCMR < 0.0003. Some studies reported the extensive metabolizer as normal metabolizer (NM) with identical cutoffs to the EM. Such data was labeled as EM.

### 2.5 Sensitivity analysis

A local sensitivity analysis of the effect of model parameters on the UCMR was performed. Individual model parameters (p_i_) were varied in both directions by 10% from the base model value 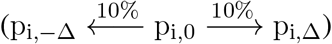 and the change in the state variable describing the UCMR at 8 hours (q) was recorded. The local sensitivity (S(q, p_i_, AS)) was calculated for a range of ASs (0, 0.25, 0.5, 1, 1.25, 1.5, 2.0, 3.0) by the following formula:

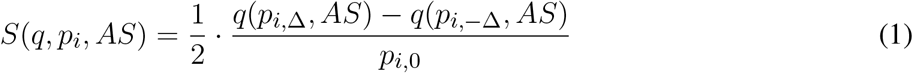

## 3 RESULTS

Within this work, a physiologically based pharmacokinetic (PBPK) model of DXM was developed and applied to study the role of the CYP2D6 polymorphism on the pharmacokinetics of DXM and metabolic phenotyping using DXM.

### 3.1 Pharmacokinetics database of DXM

For the development and evaluation of the model, a large pharmacokinetics dataset of DXM and its metabolites, consisting of 36 clinical studies, was established (Tab. 1). Most of the studies investigated either drug-gene interactions (DGI), drug-drug interactions (DDI), or the interplay of both (i.e. drug-drug-gene interactions). The large majority of studies applied DXM orally (n=35), whereas only a single publication studied DXM pharmacokinetics after intravenous application (n=1) (Duedahl et al., 2005). The application form (i.e. solution, syrup, capsule, table), the used DXM dose (2 mg to 3 mg/kg), and coadministrations (i.e. phenotyping cocktail, quinidine, cinacalcet hydrochloride, zuojin) vary between studies, as do sampling times and sampled tissues (i.e. urine, plasma, serum). Importantly, plenty of individual UCMR measurements with corresponding CYP2D6 genotype information are contained within this dataset (n=11 studies). To our knowledge, this is the first large freely available dataset of pharmacokinetics data for DXM with all data accessible from the pharmacokinetics database (PK-DB) (Grzegorzewski et al., 2021).

### 3.2 PBPK model of DXM

Within this work, a PBPK model was developed (Fig. 1) to study the role of CYP2D6 polymorphism on DXM pharmacokinetics and metabolic phenotyping with DXM. Important model parameters are provided in Tab. 2. The model is organized hierarchically, with the top layer representing the whole body (Fig. 1A) consisting of the liver, kidney, intestine, forearm, lung, and the rest compartment. Organs with minor relevance are not modeled explicitly and lumped into the rest compartment. Organs are coupled via the systemic circulation. DXM can be administered intravenously (IV) or orally (PO) with DXM appearing in the venous blood or intestine, respectively. The intestinal model (Fig 1B) describes dissolution, absorption and excretion of DXM. Only a fraction of DXM is absorbed, with the remainder excreted in the feces. DXM enters the circulatory system by crossing the enterocytes of the intestinal wall. First pass metabolism of DXM via CYP3A4 N-demethylation in the intestine reduces the amount of DXM appearing in the systemic circulation. In the liver model (Fig 1C), DXM gets transformed via O-demethylation to DXO and subsequently transformed to DXO-Glu. The reactions are modeled by Michaelis Menten kinetics and characterized with K_m_ and V_max_ values. O-demethylation takes place via CYP3A4 and CYP2D6. The K_m_ and V_max_ of CYP2D6 is modulated via the AS, details can be found in section 3.3. The kidney model (Fig 1D) describes the urinary excretion of DXM, DXO, and DXO-Glu.

**Table 2.**
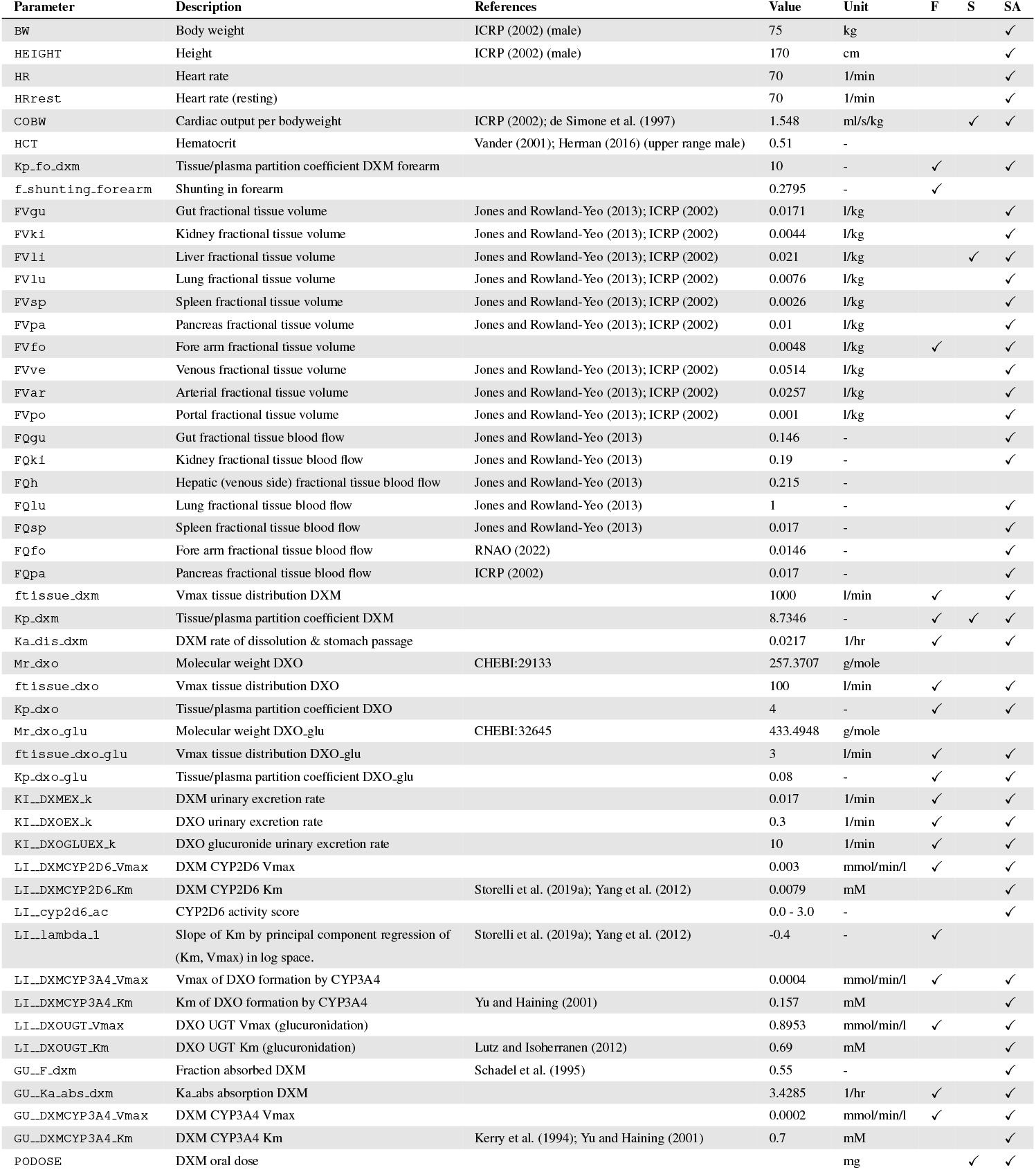
Model parameters in PBPK model of DXM. The complete information is available from the model repository. The prefixes `GU__, LI__, KI__,` correspond to the intestine/gut, liver, and kidneys, respectively. Values are either adopted from the references or fitted (**F**). During the robustness analysis of UCMR, various parameters were scanned (**S**) and a local sensitivity (**SA**) was performed, see section 3.5.

The model allows to predict concentrations and amounts of DXM, DXO, and DXO-Glu depending on CYP2D6 polymorphism, CYP2D6 diplotype, and CYP2D6 AS with amounts and concentrations of DXM, DXO, and DXO-Glu being evaluated in urine or the median cubital vein (plasma).

To our knowledge, this is the first freely accessible, reproducible, and reusable PBPK model of DXM with the model available in SBML from https://github.com/matthiaskoenig/dextromethorphan-model.

### 3.3 CYP3A4 and CYP2D6 variability

Cytochrome P450 enzymes exhibit enormous inter-individual variability in enzyme activity. To account for this variability a stochastic model of CYP2D6 and CYP3A4 activity based on bivariate lognormal distributions of K_m_ and V_max_ was developed and fitted to experimental data from human liver microsomes (Storelli et al., 2019a; Yang et al., 2012) (see Fig. 2).

For the CYP2D6 model, the V_max_ is assumed to be linearly related to the AS with *AS =* 0 having no CYP2D6 activity. The dispersion of K_m_ and V_max_ are assumed to be constant for all activity scores. For the mixture model, the frequencies of the individual activity scores P(AS) are adopted from our curated dataset (i.e. relative amount of subjects with reported activity scores and UCMRs). With increasing AS the maximal reaction velocity V_max_ of DXM conversion via CYP2D6 increases as does the affinity for the substrate DXM (K_m_ decreases). The models of CYP3A4 and CYP2D6 are capable of reproducing the data from the literature, but limited information on CYP2D6 genetics within the data hinders the validation of the AS-specific model.

As motivated in the introduction, even subjects carrying no functional variant of the CYP2D6 gene do metabolize DXM to DXO, however extremely slow. This was implemented in the model via a secondary O-demethylation via CYP3A4 with mean K_m_ for DXM adopted from Yu and Haining (2001). The dispersion of K_m_ and V_max_ is assumed to be identical to the one measured by midazolam in Storelli et al. (2019a) and Yang et al. (2012).

The resulting CYP3A4 and CYP2D6 enzyme model was coupled to the PBPK model and allowed to account (i) for the variability in DXM pharmacokinetics due to the variability in CYPs parameters and (ii) the effect of the AS on CYP2D6 activity and consequently DXM pharmacokinetics.

### 3.4 Effect of CYP2D6 activity score on DXM pharmacokinetics

Model performance was visually assessed for common pharmacokinetic measurements (i.e. DXM, DXO, DXM/DXO in plasma, and DXM/(DXO+DXO-Glu) in urine) and for subjects with reported AS or diplotype (Fig. 3). For each AS, a virtual population based on 2000 K_m_ and V_max_ samples was created from the stochastic models of CYP3A4 and CYP2D6 model. For every AS, an oral application of 30mg DXM was simulated and compared to the corresponding data. The model predicts large relative variance within a AS group as well as across different AS groups. With increasing AS, and consequently CYP2D6 activity, plasma DXM decreases (Fig. 3A), plasma DXO increases (Fig. 3B) and the plasma DXM/DXO decreases (Fig. 3C) in very good agreement with the data (Chen et al., 2017; Frank, 2009). The large variability within a AS group is a consequence of the large variability of K_m_ and V_max_ in CYP2D6 activity of a single AS (see Fig. 2). The large overlap between distributions of adjacent AS results in a large overlap in the pharmacokinetics between neighboring AS.

**Figure 3.**
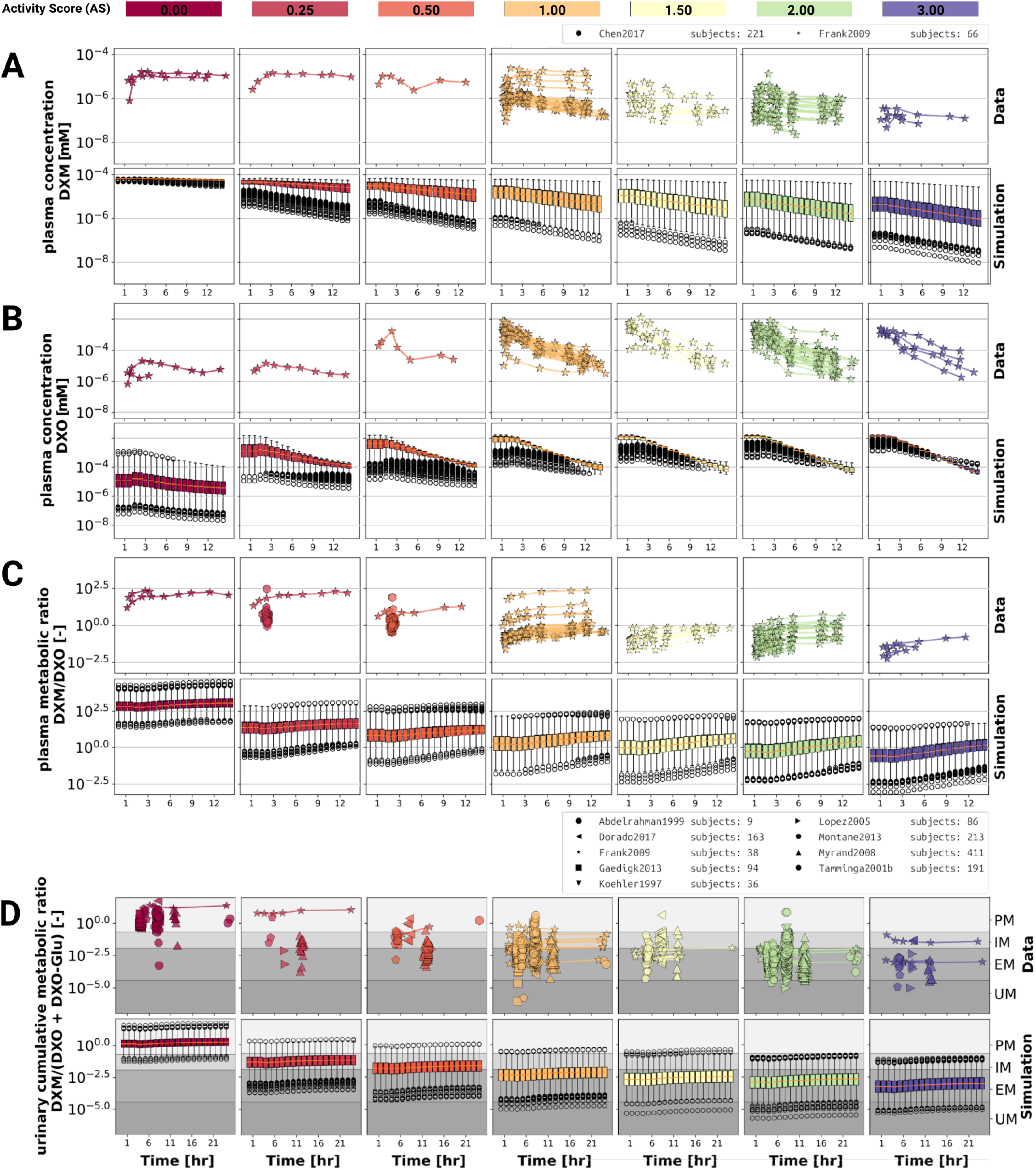
Time-dependency of DXM pharmacokinetics by activity score. **A)** DXM plasma concentration, **B)** DXO plasma concentration, **C)** DXM/DXO plasma ratio, **D**) UCMR (DXM/(DXO+DXO-Glu) in urine) Depicted is the subset of data in which 30 mg of DXM was applied orally. The upper rows in the panels depict the data in healthy adults from (Abdelrahman et al., 1999; Chen et al., 2017; Dorado et al., 2017; Frank, 2009; Gaedigk, 2013; Köhler et al., 1997; López et al., 2005; Montané Jaime et al., 2013; Myrand et al., 2008; Tamminga et al., 2001). Cocktail studies are included. Studies containing coadminstrations with established drug-drug interactions are excluded. The lower rows depict the respective simulation results. To visualize the large variability in the simulation box plots showing the quartiles along side the median and outliers for selected time points are used. Variables changed in the simulation are the CYP3A4 and CYP2D6 reaction parameters K_m_ and V_max_ according to the distributions in Fig. 2. For the different activity scores the respective CYP2D6 activity score model was used.

The UCMR (Fig. 3D) is very stable over time with a good agreement with the data. With increasing AS, the UCMR decreases and thereby shifts from PM via IM to the EM metabolic phenotype. The UCMR data was pooled independently of the amount of applied DXM (in contrast to A-C only using data from 30mg oral application) and compared to the simulation as the UCMR endpoint is very robust against the given dose (see section 3.5).

Overall the model predictions of DXM pharmacokinetics depending on AS are in very good agreement with the available data despite the limited availability of pharmacokinetics timecourses for the low AS 0, 0.25, and 0.5.

To further evaluate the model performance, simulations were compared to pharmacokinetics data for DXM in plasma or serum (Fig. 4), DXO in plasma or serum (Fig. 5), and DXO+DXO-Glu in plasma or serum (Fig. 6), DXM in urine (Fig. 7), DXO+DXO-Glu in urine (Fig. 8), and the UCMR (Fig. 9). With expected variability in mind, the model is capable to reproduce all data from the pharmacokinetics dataset. Minor shortcomings of the model are faster kinetics of DXO+DXO-Glu in plasma (Fig. 6).

**Figure 4.**
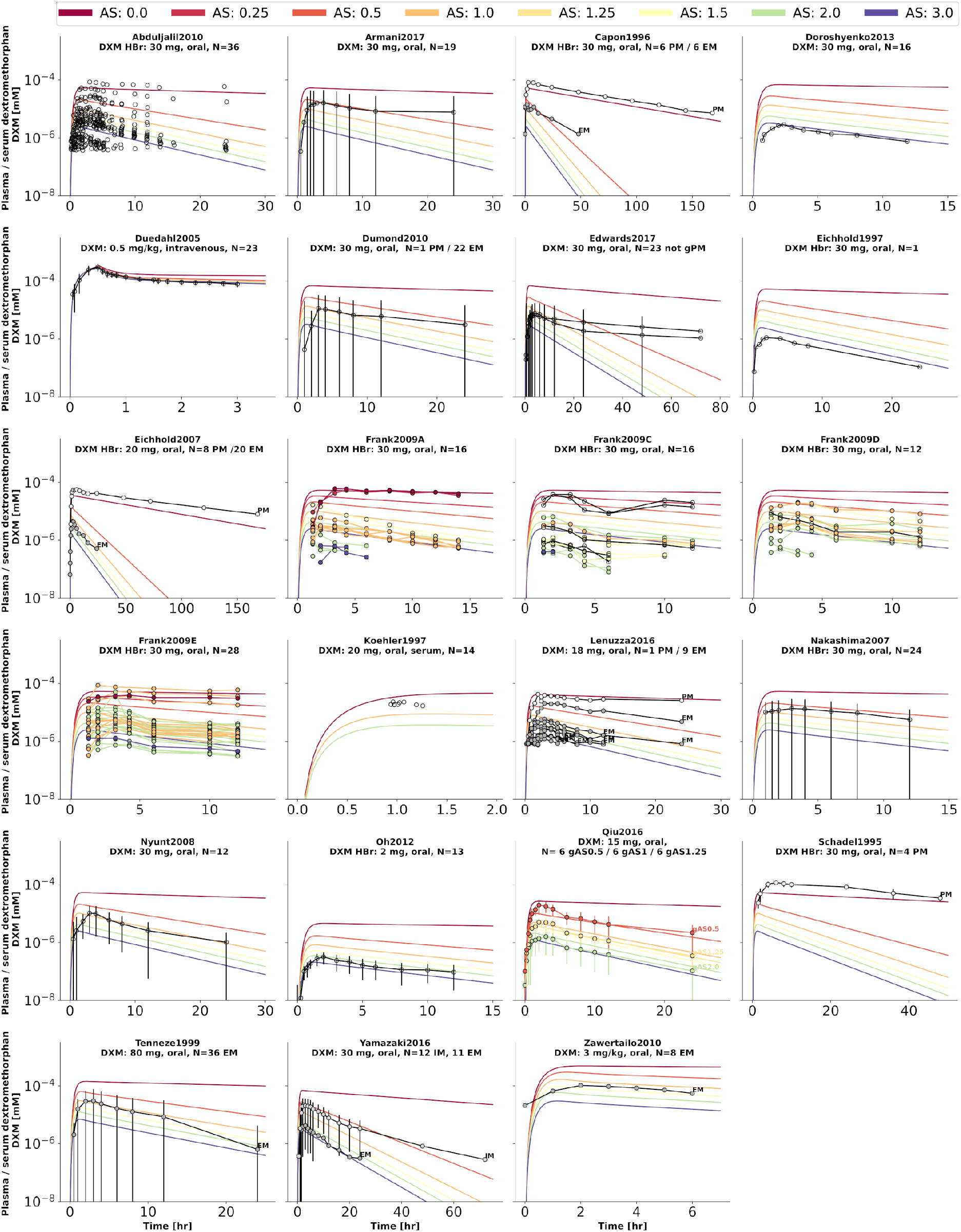
Dextromethorphan (DXM) concentration in plasma or serum. Studies were simulated according to the reported dosing protocol. In case of available activity score information the clinical data is color coded accordingly. Information on metabolizer phenotype (UM, EM, IM, PM) is provided where reported. Data from (Abduljalil et al., 2010; Armani et al., 2017; Capon et al., 1996; Doroshyenko et al., 2013; Duedahl et al., 2005; Dumond et al., 2010; Edwards et al., 2017; Eichhold et al., 1997, 2007; Frank, 2009; Köhler et al., 1997; Lenuzza et al., 2016; Nakashima et al., 2007; Nyunt et al., 2008; Oh et al., 2012; Qiu et al., 2016; Schadel et al., 1995; Tennezé et al., 1999; Yamazaki et al., 2017; Zawertailo et al., 2010).

**Figure 5.**
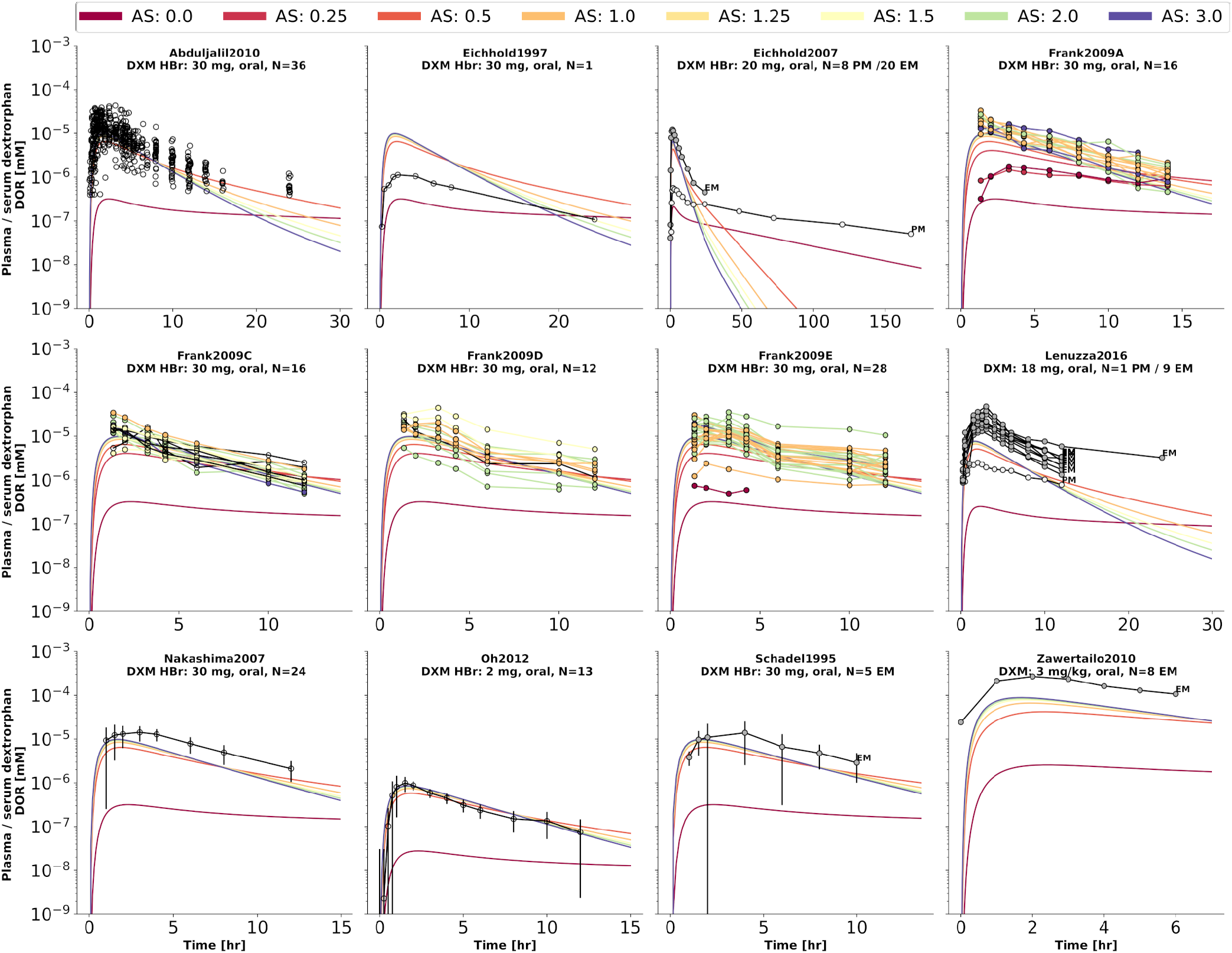
Dextrorphan (DXO) concentration in plasma or serum. Studies were simulated according to the reported dosing protocol. In case of available activity score information the clinical data is color coded accordingly. Information on metabolizer phenotype (UM, EM, IM, PM) is provided where reported. Data from (Abduljalil et al., 2010; Eichhold et al., 1997, 2007; Frank, 2009; Lenuzza et al., 2016; Nakashima et al., 2007; Oh et al., 2012; Schadel et al., 1995; Zawertailo et al., 2010).

**Figure 6.**
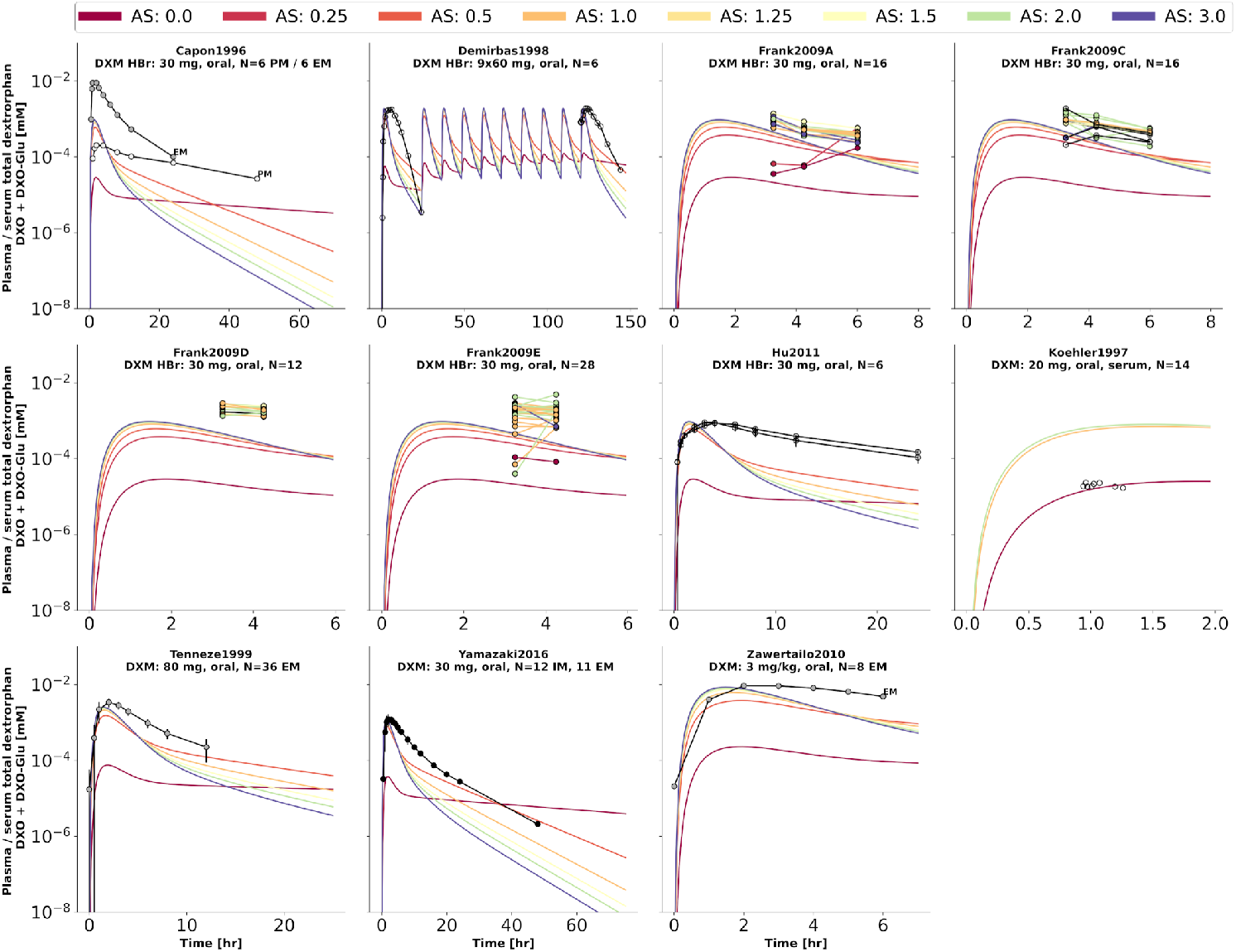
Total dextrorphan (DXO+DXO-Glu) concentration in plasma or serum. Studies were simulated according to the reported dosing protocol. In case of available activity score information the clinical data is color coded accordingly. Information on metabolizer phenotype (UM, EM, IM, PM) is provided where reported. Data from (Capon et al., 1996; Demirbas et al., 1998; Hu et al., 2011; Köhler et al., 1997; Tennezé et al., 1999; Yamazaki et al., 2017; Zawertailo et al., 2010).

**Figure 7.**
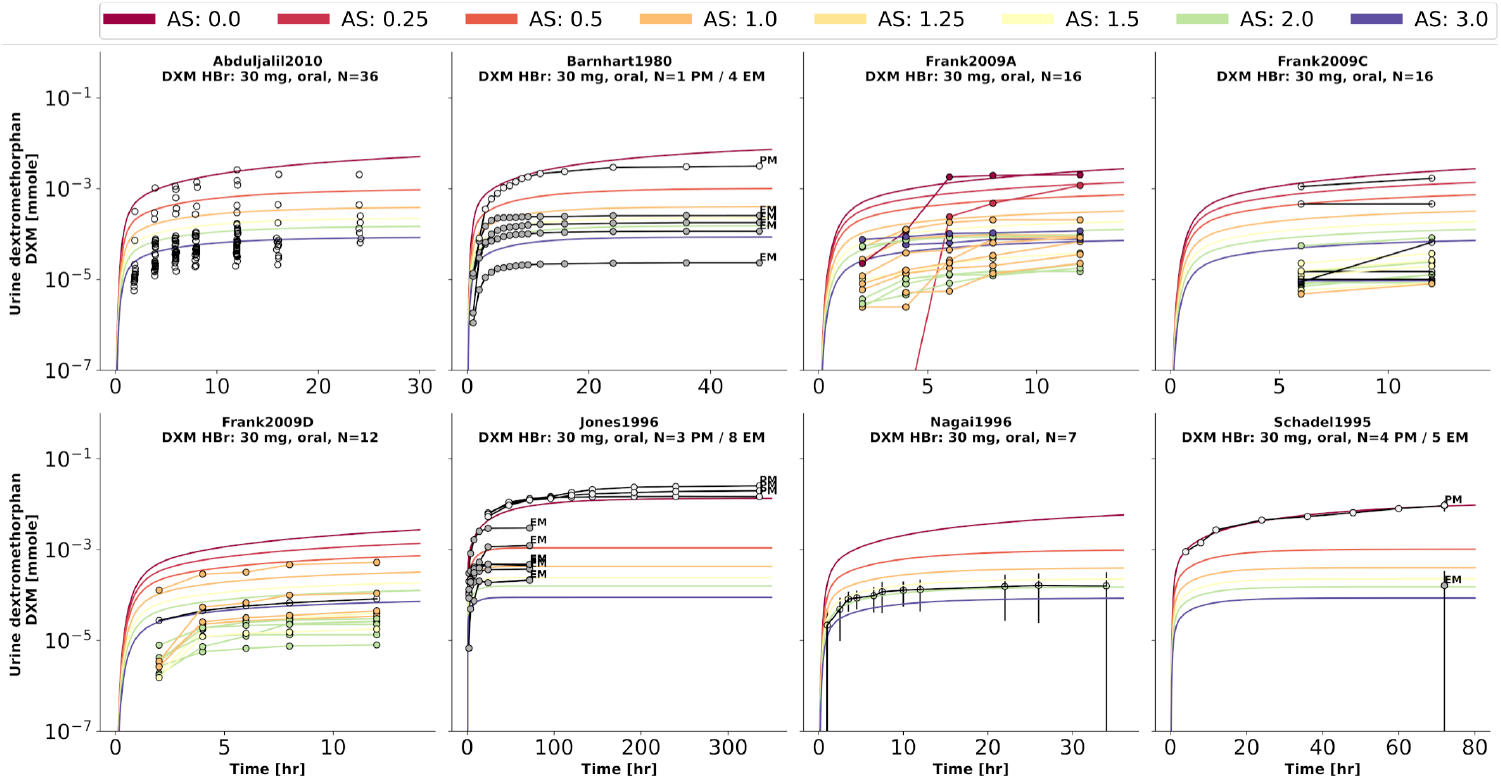
Dextromethorphan (DXM) amount in urine. Studies were simulated according to the reported dosing protocol. In case of available activity score information the clinical data is color coded accordingly. Information on metabolizer phenotype (UM, EM, IM, PM) is provided where reported. Data from (Abduljalil et al., 2010; Barnhart, 1980; Frank, 2009; Jones et al., 1996; Nagai et al., 1996; Schadel etal., 1995).

**Figure 8.**
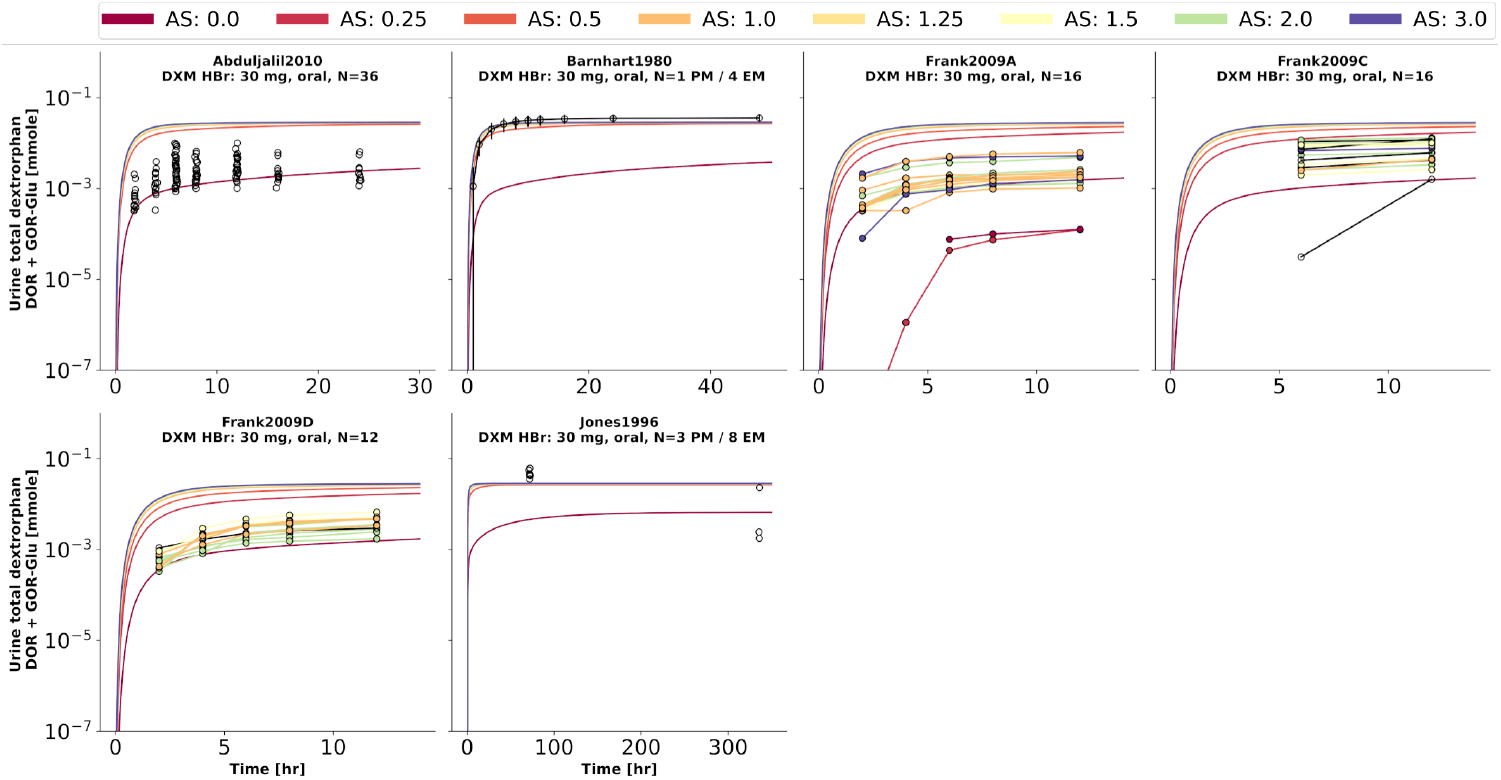
Total dextrorphan (DXO+DXO-Glu) amount in urine. Studies were simulated according to the reported dosing protocol. In case of available activity score information the clinical data is color coded accordingly. Information on metabolizer phenotype (UM, EM, IM, PM) is provided where reported. Data from (Abduljalil et al., 2010; Barnhart, 1980; Frank, 2009; Jones et al., 1996).

**Figure 9.**
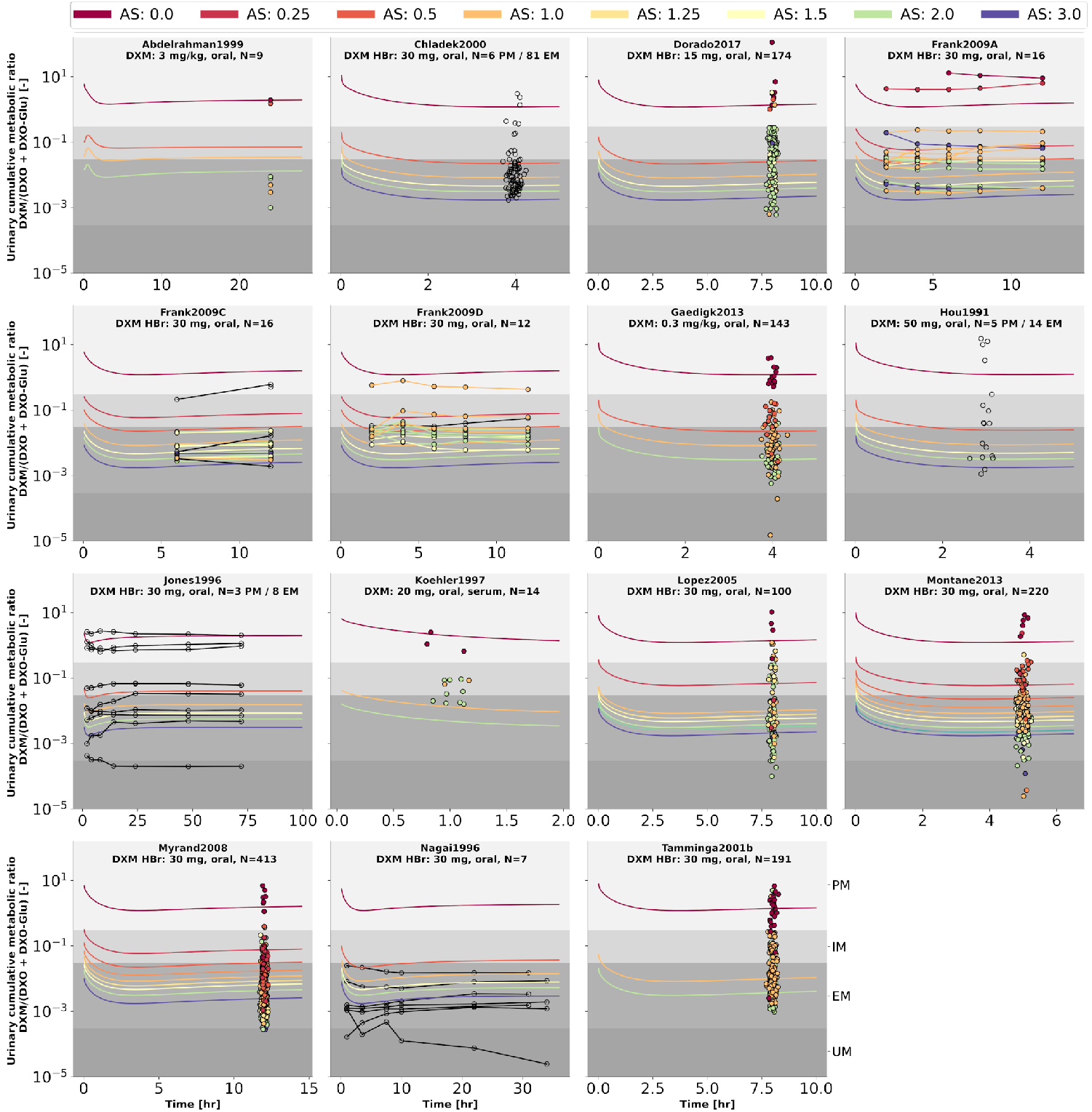
Cumulative metabolic ratio between dextormethorphan and total dextrorphan (DXM/(DXO+DXO-GLU)) in urine (UCMR). Studies were simulated according to the reported dosing protocol. In case of available activity score information the clinical data is color coded accordingly. Information on metabolizer phenotype (UM, EM, IM, PM) is provided where reported. Data from (Abdelrahman et al., 1999; Chládek et al., 2000; Dorado et al., 2017; Frank, 2009; Gaedigk, 2013; Hou et al., 1991; Jones et al., 1996; Köhler et al., 1997; López et al., 2005; Montané Jaime et al., 2013; Myrand et al., 2008; Nagai et al., 1996; Tamminga et al., 2001). The metabolic phenotype definitions for UM, EM, IM, PM are depicted as gray areas.

### 3.5 Effect of parameters on metabolic phenotyping via UCMR

Analysis of the effect of parameter changes on UCMR is highly relevant as it can help to identify potential confounding factors and bias in UCMR based phenotyping. Of special importance is the question if there is a dependency on the genetic polymorphism (activity score) of these effects.

To answer this question, model parameters (i.e. liver volume, cardiac output, tissue-to-plasma partition coefficient of DXM, and oral dose) were changed in reasonable ranges and the effect on UCMR at 8 hours after the application of 30 mg of DXM was investigated (Fig. 10A). Independent of the AS, UCMR increased with increasing liver volume and decreased with increasing cardiac output. A change in the tissue-to-plasma partition coefficient of DXM or the amount of oral DXM barely affected the UCMR.

**Figure 10.**
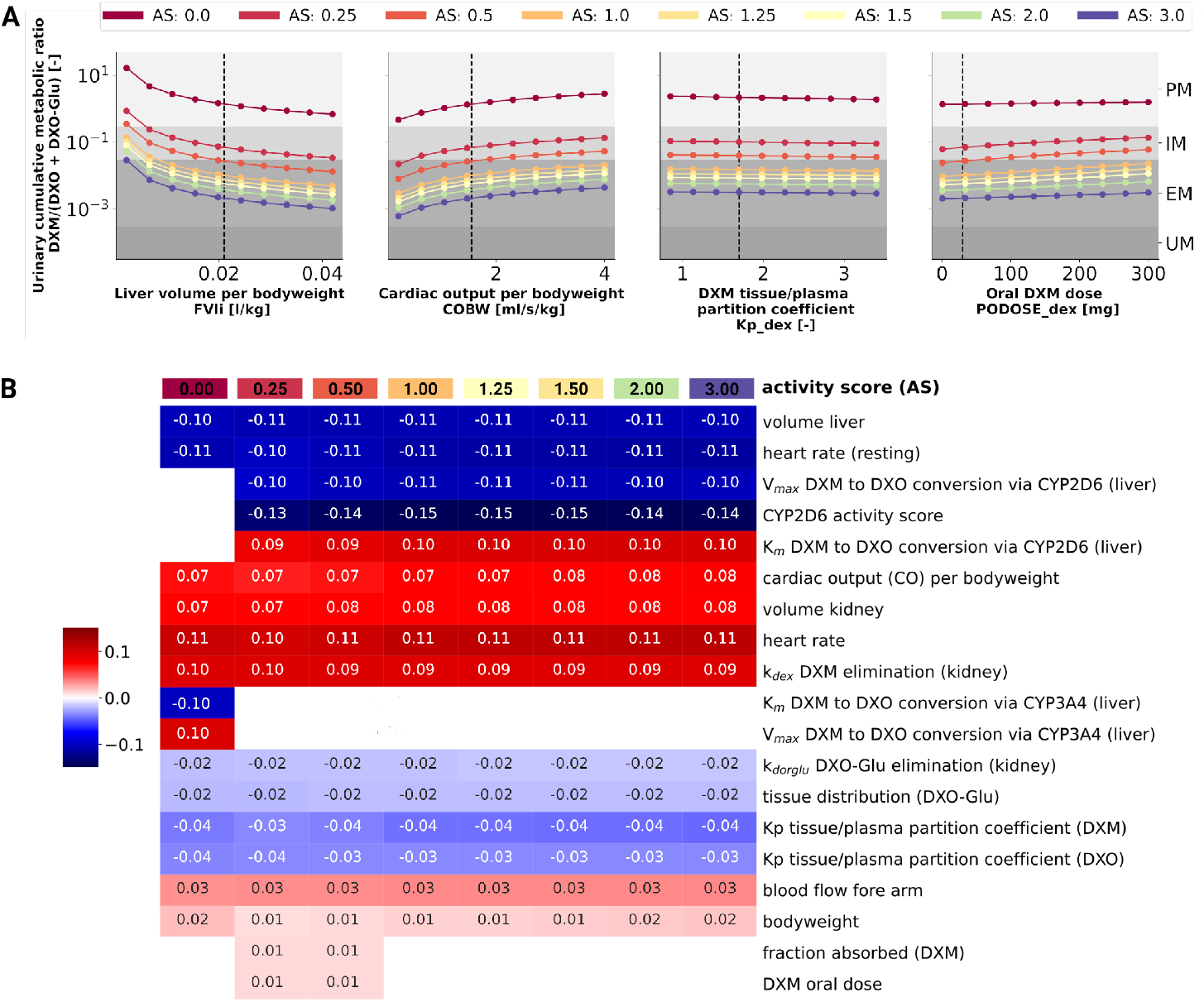
Sensitivity analysis of UCMR by activity score. **A)** Dependency of UCMR (urinary cumulative ratio of DXM/(DXO-Glu) after 8 hours and 30 mg oral DXM) on selected physiological parameters and the DXM dose. Parameter scans were performed for all activity scores. Reference model parameters are depicted as dashed lines. **B)** Sensitivity analysis of model parameters. To systematically study the effect of parameter changes the local sensitivity of UCMR were calculated for all activity scores. Parameters were varied 10% in both direction around the reference parameter value and the relative change of UCMR was calculated (insensitive parameters with relative change of UCMR smaller than 1% were omitted). Positive sensitivities are depicted in red, negative sensitivities in blue. Parameters were sorted via agglomerative clustering. Representative parameters of the clusters (i.e. liver volume per bodyweight, cardiac output per bodyweight, DXM tissue/plasma partition coefficient, and oral DXM dose) are depicted in A. The local sensitivity for the activity scores in B corresponds to the normalized slope at the dashed lines in A.

CYP2D6 phenotyping by UCMR is very stable over time as demonstrated in the time course predictions (see 3D and Fig. 9) and robust against changes in factors related to the intervention protocol (i.e. dosing amount of DXM, dissolution rate) and to some extent against changes in physiological parameters (see local sensitivity analysis of UCMR in Fig. 10B).

Liver volume, heart rate, cardiac output, kidney volume, and kidney elimination rate of DXM altered the UCMR with a similar magnitude as the CYP2D6 reaction parameters. However, the biological variation in these physiological parameters is orders of magnitude lower. The sensitivity analysis showed no effect of UGT *V_max_* and *K_m_* on UCMR which is the reason why inter-individual variability of UGT activity was not further investigated in this work. Local sensitivity of UCMR was almost identical at different AS values for almost all parameters, i.e., the effect of physiological parameters is of similar relative magnitude independent of AS. For AS=0, our model assumptions of minor DXM metabolism by CYP3A4 lead to UCMR not being modulated by CYP2D6 but rather by CYP3A4 activity. Nonetheless, even across studies with non-standardized intervention protocols the UCMR measurements seams to be a good but not perfect endpoint to quantify and compare CYP2D6 enzyme activity. Importantly, our analysis indicates that UCMR measurements can be pooled even across investigations with different intervention protocols (as for instance performed in Fig. 3D). This still may lead to biases and errors, e.g. due to differences in the quantification protocol.

### 3.6 Effect of CYP2D6 polymorphisms and activity score on UCMR

Next, we tested if the model is able to predict UCMR distributions for given genotypes and AS (Fig. 11). Model predictions based on underlying genotype frequencies were compared with the experimental data. UCMR distributions for individual AS groups are well described by the model. The AS impacts the UCMR, with increasing AS resulting in an decrease in UCMR. However individual AS distributions heavily overlap, as expected, due to the large overlap in CYP2D6 parameter distributions between different AS. The predicted distributions tend to be slightly narrower than the actual data. Possible reasons are many fold (e.g. omitted physiological variation, omitted variation in UGT activity, difficulties in correct genotype assignment, unknown effect modifiers, and biases).

**Figure 11.**
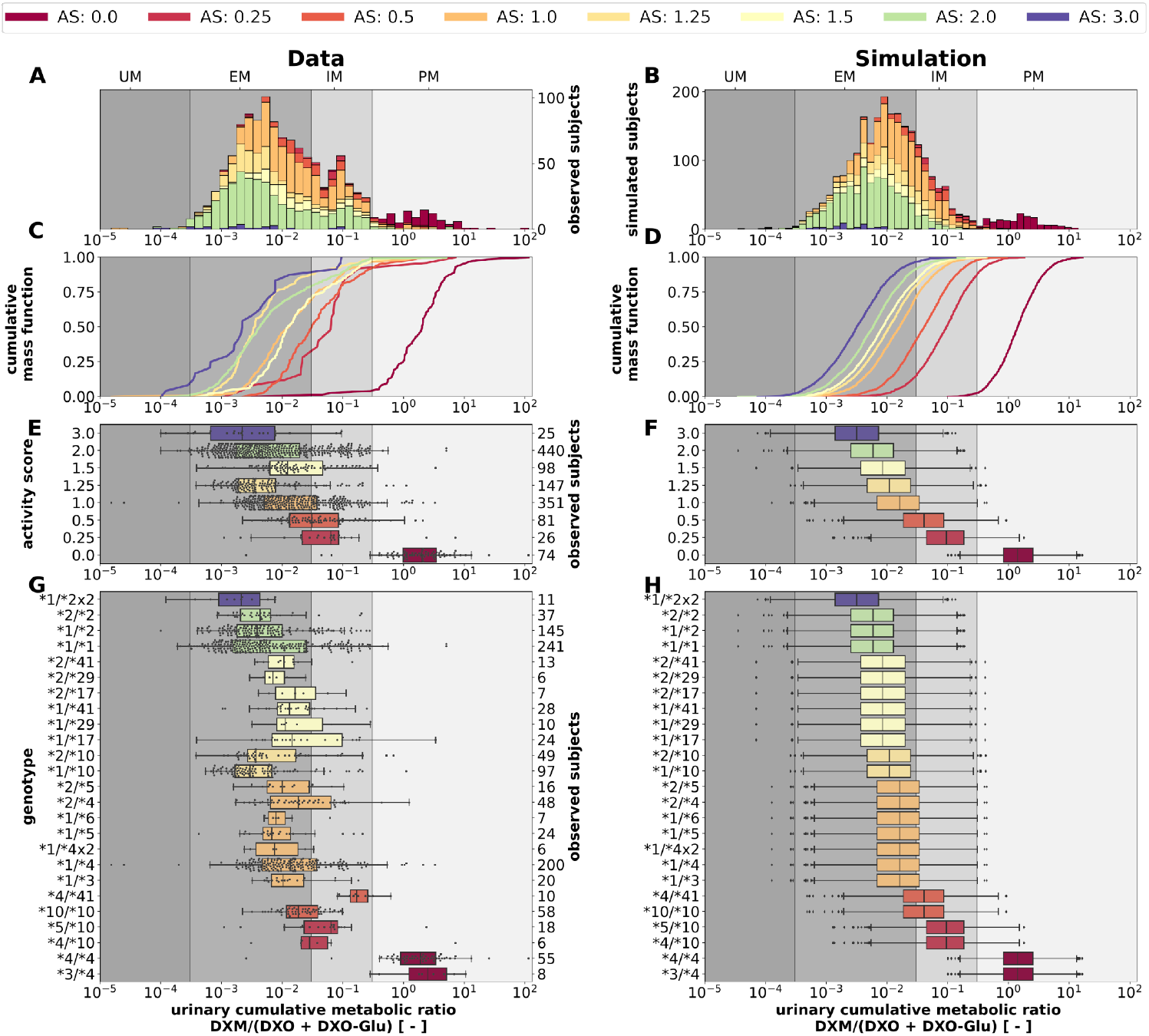
CYP2D6 genotype-, activity score association of the UCMR. Simulation of urinary cumulative ratio of DXM/(DXO-Glu) (UCMR) based on activity score frequencies. UCMR data was measured at least 4 hours after the application of DXM (hydrobromide) in healthy adults. Cocktail studies were included in the analysis. Studies containing coadminstrations with established drug-drug interactions were excluded. The ranges for metabolic phenotypes (UM, EM, IM, PM) are depicted as gray shaded areas. For timecourse UCMRs, only the latest measurement after administration was included. Data from (Abdelrahman et al., 1999; Dorado et al., 2017; Frank, 2009; Gaedigk, 2013; Köhler et al., 1997; López et al., 2005; Montané Jaime et al., 2013; Myrand et al., 2008; Tamminga et al., 2001). **A)** Histogram of UCMR data stratified by CYP2D6 activity score. **B)** Corresponding simulation results (**UCMR** at 8hr) from the Monte Carlo simulation with random variables being the enzyme reaction parameter (i.e. K_m_, V_max_). See details in Fig. 2. **C)** Empirical CMFs stratified by the activity scores. **D)** Corresponding simulated CMFs stratified by CYP2D6 activity scores. **E)** Box plots of observed UCMRs stratified by CYP2D6 activity scores. **F)** Box plots of simulated UCMRs stratified by the activity scores. **G)** Box plots of observed UCMRs stratified by CYP2D6 diplotypes. **H)** Box plots of simulated UCMRs stratified by CYP2D6 diplotypes. For D, F, and H, 2000 samples were simulated for each activity score whereas in B and D a two-fold oversampling with the CYP2D6 activity score frequencies from the UCMR data was performed.

The AS system could be refined to better describe the data. The categorization of CYP2D6 genotypes into discrete activity values (i. e. 0, 0.25, 0.5, 1) is an oversimplification, a continuous activity score would probably perform better. The model and data indicate that gUM (AS ≥ 3) is a very unreliable predictor for ultra rapid metabolism and only gPMs (AS = 0) are almost perfectly distinguishable from other metabolizers, see Fig. 11C, D, E, and F.

Another strength of the presented model is that it enables the prediction of the *in vivo* phenotype of subjects based on *in vitro* data.

### 3.7 Population variability in UCMR

Finally, the model was also capable to predict UCMR distributions for different biogeographical populations (Fig. 12) based on the underlying AS frequencies (Tab. S2). Based on the reported frequencies, the UCMR distributions were simulated at 8 hours after the application of 30mg DXM for Oceanian, Near Eastern, American, Latino, Central/South Asian, African American/Afro-Caribbean, Sub-Saharan African, European, and East Asian populations (Fig. 12A). Data for Caucasian and East Asian populations (Fig. 12B) was used for validation of the predictions (Fig. 12C). The data is in good agreement with measurements of Caucasians and East Asians as reported by Abdelrahman et al. (1999); Frank (2009); Gaedigk (2013); Köhler et al. (1997); Myrand et al. (2008).

**Figure 12.**
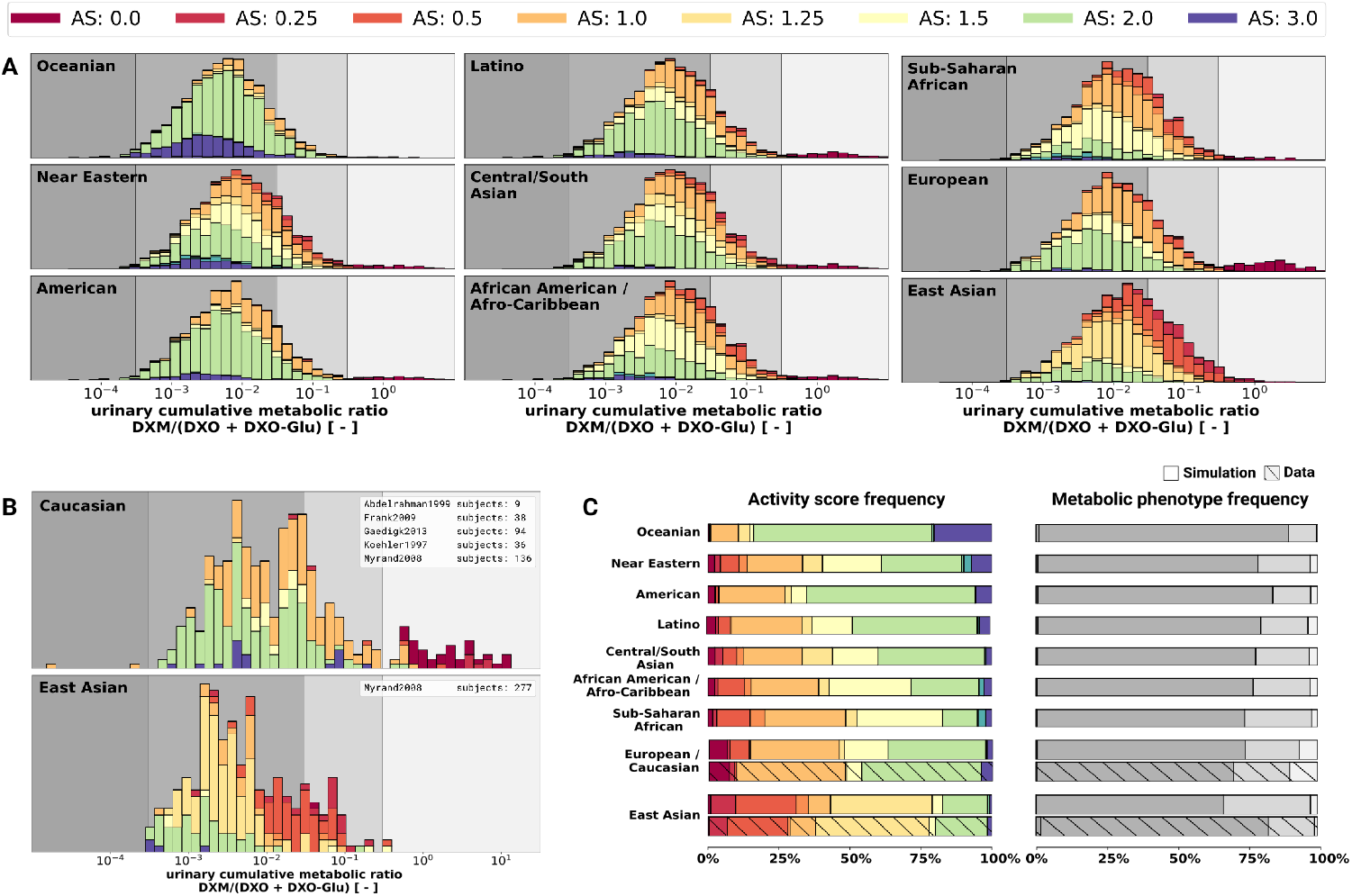
UCMR distributions for biogeographical populations. **A)** Simulated UCMR distributions at 8 hours for various biogeographical populations based on reported CYP2D6 activity score frequencies as reported in PharmGKB (Whirl-Carrillo et al., 2021). Frequencies are provided in the Supplementary Tab. S2. **B)** Reported UCMRs depending on activity score for Caucasians and East Asians from (Abdelrahman et al., 1999; Frank, 2009; Gaedigk, 2013; Köhler et al., 1997; Myrand et al., 2008). Cocktail studies were included in the analysis. Studies containing coadminstrations with established drug-drug interactions were excluded. **C)** Simulated activity score frequency and metabolic phenotype frequency for the populations and comparison with data for Caucasian and East Asian populations (hatched bars).

## 4 DISCUSSION

During the last 20 years various modeling approaches and software solutions were utilized to investigate various aspects of DXM pharmacokinetics, e.g. using GastroPlus (Bolger et al., 2019), P-Pharm (Moghadamnia et al., 2003), SAS (Ito et al., 2010; Chiba et al., 2012), SimCYP (Dickinson et al., 2007; Ke et al., 2013; Sager et al., 2014; Chen et al., 2016; Rougée et al., 2016; Adiwidjaja et al., 2018; Machavaram et al., 2019; Storelli et al., 2019b), MATLAB (Kim et al., 2017), or PK-Sim (Rüdesheim et al., 2022). However, most of the work is difficult/impossible to validate or to build up on due to a lack of accessibility of models and software, and platform-dependency of the models. Here, we provide an openly accessible, reproducible and platform-independent whole-body model of DXM metabolism, which facilitates reusability, extensibility, and comparability.

Apart from that, modeling work which aims for high empirically evidence relies on trustworthy supporting real world data. More and independent sources of data are highly beneficial for the scientific outcomes. For that matter, guidelines like PRISMA for reporting transparency, completeness, and accuracy find very broad endorsement in the field of systematic reviews and meta analysis. The present work faces somewhat similar challenges for the evaluation and selection of data from literature. Therefore, PRISMA-ScR guidelines were adopted where applicable. With this approach, bias within the used dataset could be mitigated or at least identified. Importantly, we supplement our open and accessible model with a large, open, and accessible database of pharmacokinetics data.

The presented PBPK model is able to predict the DXM metabolism of populations and individuals based on their CYP2D6 genotype. It is probably the first model capable to predict individual UCMRs and the expected distributions of UCMR. Moreover, it can reproduce a broad range of reported clinical data on DXM and enables better intuition on how to interpret DXM related pharmacokinetics. E.g., an important message is that CYP2D6 activity is not the only modulator of UCMR, as can be seen by the large variability in activity score and overlap between activity scores. UCMR as a proxy of CYP2D6 metabolic phenotype should therefore be interpreted carefully. The model shows that for extremely low CYP2D6 activity the UCMR is not primarily governed by the CYP2D6 activity. This is consistent with the finding that CYP2D6 inhibition merely affects PMs (Pope et al., 2004).

The current version of the model is already very valuable, still there is plenty of room for improvement. By providing the data and model in open and standardized formats we enable and encourage these improvements by model extensions and updates.

Many of the physiological parameters in the model were fitted or estimated although they could be measured in principle. E.g., relatively low DXM concentrations in plasma suggest substantial extra-vascular binding of DXM. However, tissue-plasma partition coefficients (K_p_) are difficult to assess and only limited data is available. Steinberg et al. (1996) reported brain levels to be 68-fold higher and cerebrospinal fluid levels 4-fold lower than serum levels, respectively. Others estimated K_p_ ~ 1.65 from n-octanol-water partition coefficients and again others suggested additional trapping mechanisms (i.e. lysosomal trapping) (Bolger et al., 2019). In the model, the DXO-Glu kinetics is a bit to rapid (see Fig. 6), probably due to the decision to model tissue distribution uniformly for all organs (i.e. identical K_p_ and f_tissue_). We decided for a more parsimonious model. Glucuronides, however, are generally much more polar than their respective non-glucuronides which result in less plasma binding, higher urinary excretion, lower lipid-solubility, and higher water-solubility. Transport into different tissues is affected differently by polarity.

Most important for model improvements would be additional *in vitro* measurements on the association between CYP2D6 genotype and phenotype which are very limited in literature (Storelli et al., 2019a; Ning et al., 2019; Dalton et al., 2020). Furthermore, simultaneous in *vitro* and UCMR measurements do not exist the literature. Both would be very important for the validation of the AS system and the development of new models which e.g. take into account structural variation (Dalton et al., 2020). For instance, with the AS system alone it is not possible to explain why CYP2D6 is inhibited differently for different genotypes Qiu et al. (2016).

In conclusion, we developed and validated a PBPK model of DXM and applied it to study the effect of the CYP2D6 polymorphism on metabolic phenotyping.

## AUTHOR CONTRIBUTIONS

JG and MK designed the study, developed the computational model, implemented and performed the analysis, and wrote the initial draft of the manuscript. JB provided support with data curation. All authors discussed the results. All authors contributed to and revised the manuscript critically.

## DATA AVAILABILITY STATEMENT

All clinical data of dextromethorphan pharmacokinetics that was used in this work can be found in PK-DB available from https://pk-db.com.

## REFERENCES

Abdelrahman, S., Gotschall, R., Kauffman, R., Leeder, J., and Kearns, G. (1999). Investigation of terbinafine as a CYP2D6 inhibitor in vivo. Clinical Pharmacology & Therapeutics 65, 465–472. doi:10.1016/S0009-9236(99)70065-2

Abduljalil, K., Frank, D., Gaedigk, A., Klaassen, T., Tomalik-Scharte, D., Jetter, A., et al. (2010). Assessment of Activity Levels for CYP2D6*1, CYP2D6*2, and CYP2D6*41 Genes by Population Pharmacokinetics of Dextromethorphan. Clinical Pharmacology & Therapeutics 88, 643–651. doi:10.1038/clpt.2010.137

Adiwidjaja, J., Boddy, A. V., and McLachlan, A. J. (2018). A Strategy to Refine the Phenotyping Approach and Its Implementation to Predict Drug Clearance: A Physiologically Based Pharmacokinetic Simulation Study. CPT: Pharmacometrics & Systems Pharmacology 7, 798–808. doi:10.1002/psp4.12355

Armani, S., Ting, L., Sauter, N., Darstein, C., Tripathi, A. P., Wang, L., et al. (2017). Drug Interaction Potential of Osilodrostat (LCI699) Based on Its Effect on the Pharmacokinetics of Probe Drugs of Cytochrome P450 Enzymes in Healthy Adults. Clinical drug investigation 37, 465–472. doi:10.1007/s40261-017-0497-0

Barnhart, J. W. (1980). The urinary excretion of dextromethorphan and three metabolites in dogs and humans. Toxicology and Applied Pharmacology 55, 43–48. doi:10.1016/0041-008X(80)90218-5

Berm, E. J. J., Risselada, A. J., Mulder, H., Hak, E., and Wilffert, B. (2013). Phenoconversion of cytochrome P450 2D6: The need for identifying the intermediate metabolizer genotype. The Journal of clinical psychiatry 74, 1025. doi:10.4088/JCP.13lr08555

Bolger, M. B., Macwan, J. S., Sarfraz, M., Almukainzi, M., and Löbenberg, R. (2019). The Irrelevance of In Vitro Dissolution in Setting Product Specifications for Drugs Like Dextromethorphan That are Subject to Lysosomal Trapping. Journal of pharmaceutical sciences 108, 268–278. doi:10.1016/j.xphs.2018.09.036

Capon, D. A., Bochner, F., Kerry, N., Mikus, G., Danz, C., and Somogyi, A. A. (1996). The influence of CYP2D6 polymorphism and quinidine on the disposition and antitussive effect of dextromethorphan in humans. Clinical Pharmacology & Therapeutics 60, 295–307. doi:10.1016/S0009-9236(96)90056-9

Caudle, K. E., Sangkuhl, K., Whirl-Carrillo, M., Swen, J. J., Haidar, C. E., Klein, T. E., et al. (2020). Standardizing CYP2D6 Genotype to Phenotype translation: Consensus Recommendations from the Clinical Pharmacogenetics Implementation Consortium and Dutch Pharmacogenetics Working Group. Clinical and Translational Science 13, 116–124. doi:10.1111/cts.12692

Chen, R., Rostami-Hodjegan, A., Wang, H., Berk, D., Shi, J., and Hu, P. (2016). Application of a physiologically based pharmacokinetic model for the evaluation of single-point plasma phenotyping method of CYP2D6. European journal of pharmaceutical sciences: official journal of the European Federation for Pharmaceutical Sciences 92, 131–136. doi:10.1016/j.ejps.2016.07.001

Chen, R., Zheng, X., and Hu, P. (2017). CYP2D6 Phenotyping Using Urine, Plasma, and Saliva Metabolic Ratios to Assess the Impact of CYP2D6_*_ 10 on Interindividual Variation in a Chinese Population. Frontiers in Pharmacology 8, 239. doi:10.3389/fphar.2017.00239

Chiba, K., Kato, M., Ito, T., Suwa, T., and Sugiyama, Y. (2012). Inter-individual variability of in vivo CYP2D6 activity in different genotypes. Drug metabolism and pharmacokinetics 27, 405–413. doi:10.2133/dmpk.dmpk-11-rg-078

Chládek, J., Zimová, G., Beránek, M., and Martínková, J. (2000). In-vivo indices of CYP2D6 activity: Comparison of dextromethorphan metabolic ratios in 4-h urine and 3-h plasma. European Journal of Clinical Pharmacology 56, 651–657. doi:10.1007/s002280000218

Dalton, R., Lee, S.-B., Claw, K. G., Prasad, B., Phillips, B. R., Shen, D. D., et al. (2020). Interrogation of CYP2D6 Structural Variant Alleles Improves the Correlation Between CYP2D6 Genotype and CYP2D6-Mediated Metabolic Activity. Clinical and translational science 13, 147–156. doi:10.1111/cts.12695

De Kesel, P. M. M., Lambert, W. E., and Stove, C. P. (2016). Alternative Sampling Strategies for Cytochrome P450 Phenotyping. Clinical Pharmacokinetics 55, 169–184. doi:10.1007/s40262-015-0306-y

de Simone, G., Devereux, R. B., Daniels, S. R., Mureddu, G., Roman, M. J., Kimball, T. R., et al. (1997). Stroke volume and cardiac output in normotensive children and adults. Assessment of relations with body size and impact of overweight. Circulation 95, 1837–1843. doi:10.1161/01.cir.95.7.1837

Demirbas, S., Reyderman, L., and Stavchansky, S. (1998). Bioavailability of dextromethorphan (as dextrorphan) from sustained release formulations in the presence of guaifenesin in human volunteers. Biopharmaceutics & Drug Disposition 19, 541–545. doi:10.1002/(SICI)1099-081X(1998110)19:8〈541::AID-BDD138〉3.0.CO;2-8

Dickinson, G. L., Rezaee, S., Proctor, N. J., Lennard, M. S., Tucker, G. T., and Rostami-Hodjegan, A. (2007). Incorporating In Vitro Information on Drug Metabolism Into Clinical Trial Simulations to Assess the Effect of CYP2D6 Polymorphism on Pharmacokinetics and Pharmacodynamics: Dextromethorphan as a Model Application. The Journal of Clinical Pharmacology 47, 175–186. doi:10.1177/0091270006294279

Dorado, P., González, I., Naranjo, M. E. G., de Andrés, F., Peñas-Lledó, E. M., Calzadilla, L. R., et al. (2017). Lessons from Cuba for Global Precision Medicine: CYP2D6 Genotype Is Not a Robust Predictor of CYP2D6 Ultrarapid Metabolism. OMICS: A Journal of Integrative Biology 21, 17–26. doi:10.1089/omi.2016.0166

Doroshyenko, O., Rokitta, D., Zadoyan, G., Klement, S., Schläfke, S., Dienel, A., et al. (2013). Drug cocktail interaction study on the effect of the orally administered lavender oil preparation silexan on cytochrome P450 enzymes in healthy volunteers. Drug Metab. Dispos. 41, 987–993. doi:10.1124/dmd.112.050203

Duedahl, T. H., Dirks, J., Petersen, K. B., Romsing, J., Larsen, N.-E., and Dahl, J. B. (2005). Intravenous dextromethorphan to human volunteers: Relationship between pharmacokinetics and anti-hyperalgesic effect. Pain 113, 360–368. doi:10.1016/j.pain.2004.11.015

Dumond, J. B., Vourvahis, M., Rezk, N. L., Patterson, K. B., Tien, H.-C., White, N., et al. (2010). A Phenotype–Genotype Approach to Predicting CYP450 and P-Glycoprotein Drug Interactions With the Mixed Inhibitor/Inducer Tipranavir/Ritonavir. Clinical Pharmacology & Therapeutics 87, 735–742. doi:10.1038/clpt.2009.253

Edwards, J. E., Eliot, L., Parkinson, A., Karan, S., and MacConell, L. (2017). Assessment of Pharmacokinetic Interactions Between Obeticholic Acid and Caffeine, Midazolam, Warfarin, Dextromethorphan, Omeprazole, Rosuvastatin, and Digoxin in Phase 1 Studies in Healthy Subjects. Advances in Therapy 34, 2120–2138. doi:10.1007/s12325-017-0601-0

Eichelbaum, M., Spannbrucker, N., Steincke, B., and Dengler, H. J. (1979). Defective N-oxidation of sparteine in man: A new pharmacogenetic defect. European journal of clinical pharmacology 16, 183–187. doi:10.1007/BF00562059

Eichhold, T. H., Greenfield, L. J., Hoke, S. H., and Wehmeyer, K. R. (1997). Determination of dextromethorphan and dextrorphan in human plasma by liquid chromatography/tandem mass spectrometry. Journal of Mass Spectrometry 32, 1205–1211. doi:10.1002/(SICI)1096-9888(199711)32:11〈1205::AID-JMS579〉3.0.CO;2-C

Eichhold, T. H., McCauley-Myers, D. L., Khambe, D. A., Thompson, G. A., and Hoke, S. H. (2007). Simultaneous determination of dextromethorphan, dextrorphan, and guaifenesin in human plasma using semi-automated liquid/liquid extraction and gradient liquid chromatography tandem mass spectrometry. Journal of Pharmaceutical and Biomedical Analysis 43, 586–600. doi:10.1016/j.jpba.2006.07.018

Frank, D. (2009). Evaluation of pharmacokinetic metrics for phenotyping of the human CYP2D6 enzyme with dextromethorphan. Ph.D. thesis, Rheinische Friedrich-Wilhelms-Universität Bonn

Frank, D., Jaehde, U., and Fuhr, U. (2007). Evaluation of probe drugs and pharmacokinetic metrics for CYP2D6 phenotyping. European Journal of Clinical Pharmacology 63, 321–333. doi:10.1007/s00228-006-0250-8

Fuhr, U., Jetter, A., and Kirchheiner, J. (2007). Appropriate phenotyping procedures for drug metabolizing enzymes and transporters in humans and their simultaneous use in the “cocktail” approach. Clinical pharmacology and therapeutics 81, 270–283. doi:10.1038/sj.clpt.6100050

Gaedigk, A. (2013). Complexities of CYP2D6 gene analysis and interpretation. International Review of Psychiatry 25, 534–553. doi:10.3109/09540261.2013.825581

Gaedigk, A., Dinh, J., Jeong, H., Prasad, B., and Leeder, J. (2018a). Ten Years’ Experience with the CYP2D6 Activity Score: A Perspective on Future Investigations to Improve Clinical Predictions for Precision Therapeutics. Journal of Personalized Medicine 8, 15. doi:10.3390/jpm8020015

Gaedigk, A., Eklund, J. D., Pearce, R. E., Leeder, J. S., Alander, S. W., Phillips, M. S., et al. (2007). Identification and characterization of CYP2D6*56B, an allele associated with the poor metabolizer phenotype. Clinical pharmacology and therapeutics 81, 817–820. doi:10.1038/sj.clpt.6100125

Gaedigk, A., Ingelman-Sundberg, M., Miller, N. A., Leeder, J. S., Whirl-Carrillo, M., Klein, T. E., et al. (2018b). The pharmacogene variation (PharmVar) consortium: Incorporation of the human cytochrome P450 (CYP) allele nomenclature database. Clinical pharmacology and therapeutics 103, 399–401. doi:10.1002/cpt.910

Gaedigk, A., Sangkuhl, K., Whirl-Carrillo, M., Klein, T., and Leeder, J. S. (2017). Prediction of CYP2D6 phenotype from genotype across world populations. Genetics in Medicine: Official Journal of the American College of Medical Genetics 19, 69–76. doi:10.1038/gim.2016.80

Gaedigk, A., Turner, A., Everts, R. E., Scott, S. A., Aggarwal, P., Broeckel, U., et al. (2019). Characterization of Reference Materials for Genetic Testing of CYP2D6 Alleles. The Journal of Molecular Diagnostics 21, 1034–1052. doi:10.1016/j.jmoldx.2019.06.007

Gasche, Y., Daali, Y., Fathi, M., Chiappe, A., Cottini, S., Dayer, P., et al. (2004). Codeine intoxication associated with ultrarapid CYP2D6 metabolism. The New England journal of medicine 351, 2827–2831. doi:10.1056/NEJMoa041888

Gonzalez Hernandez, F., Carter, S. J., Iso-Sipilä, J., Goldsmith, P., Almousa, A. A., Gastine, S., et al. (2021). An automated approach to identify scientific publications reporting pharmacokinetic parameters. Wellcome Open Research 6, 88. doi:10.12688/wellcomeopenres.16718.1

Grzegorzewski, J., Bartsch, F., Köller, A., and König, M. (2022). Pharmacokinetics of Caffeine: A Systematic Analysis of Reported Data for Application in Metabolic Phenotyping and Liver Function Testing. Frontiers in Pharmacology 12. doi:10.3389/fphar.2021.752826

Grzegorzewski, J., Brandhorst, J., Green, K., Eleftheriadou, D., Duport, Y., Barthorscht, F., et al. (2021). PK-DB: Pharmacokinetics database for individualized and stratified computational modeling. Nucleic Acids Research 49, D1358–D1364. doi:10.1093/nar/gkaa990

[Dataset] Grzegorzewski, J. and König, M. (2022). Physiologically based pharmacokinetic (PBPK) model of dextromethorphan. https://doi.org/10.5281/zenodo.7025683. doi:10.5281/zenodo.7025683. Accessed: 2022-08-26

Herman, I. P. (2016). Physics of the Human Body (Springer)

Hou, Z.-Y., Pickle, L. W., Meyer, P. S., and Woosley, R. L. (1991). Salivary analysis for determination of dextromethorphan metabolic phenotype. Clinical Pharmacology and Therapeutics 49, 410–419. doi:10.1038/clpt.1991.48

Hu, L., Li, L., Yang, X., Liu, W., Yang, J., Jia, Y., et al. (2011). Floating matrix dosage form for dextromethorphan hydrobromide based on gas forming technique: In vitro and in vivo evaluation in healthy volunteers. European Journal of Pharmaceutical Sciences 42, 99–105. doi:10.1016/j.ejps.2010.10.010

Hucka, M., Bergmann, F. T., Chaouiya, C., Dräger, A., Hoops, S., Keating, S. M., et al. (2019). The systems biology markup language (SBML): Language specification for level 3 version 2 core release 2. Journal of Integrative Bioinformatics doi:10.1515/jib-2019-0021

Hurtado, I., García-Sempere, A., Peiró, S., and Sanfélix-Gimeno, G. (2020). Increasing Trends in Opioid Use From 2010 to 2018 in the Region of Valencia, Spain: A Real-World, Population-Based Study. Frontiers in Pharmacology 11, 612556. doi:10.3389/fphar.2020.612556

ICRP (2002). Basic anatomical and physiological data for use in radiological protection: Reference values. A report of age-and gender-related differences in the anatomical and physiological characteristics of reference individuals. ICRP Publication 89. Annals of the ICRP 32, 5–265

Ito, T., Kato, M., Chiba, K., Okazaki, O., and Sugiyama, Y. (2010). Estimation of the interindividual variability of cytochrome 2D6 activity from urinary metabolic ratios in the literature. Drug Metabolism and Pharmacokinetics 25, 243–253. doi:10.2133/dmpk.25.243

Jones, D. R., Gorski, J. C., Haehner, B. D., O’Mara, E. M., and Hall, S. D. (1996). Determination of cytochrome P450 3A4/5 activity in vivo with dextromethorphan N-demethylation. Clinical Pharmacology and Therapeutics 60, 374–384. doi:10.1016/S0009-9236(96)90194-0

Jones, H. and Rowland-Yeo, K. (2013). Basic concepts in physiologically based pharmacokinetic modeling in drug discovery and development. CPT: pharmacometrics & systems pharmacology 2, e63. doi:10.1038/psp.2013.41

Kawanishi, C., Lundgren, S., Agren, H., and Bertilsson, L. (2004). Increased incidence of CYP2D6 gene duplication in patients with persistent mood disorders: Ultrarapid metabolism of antidepressants as a cause of nonresponse. A pilot study. European Journal of Clinical Pharmacology 59, 803–807. doi:10.1007/s00228-003-0701-4

Ke, A. B., Nallani, S. C., Zhao, P., Rostami-Hodjegan, A., Isoherranen, N., and Unadkat, J. D. (2013). A Physiologically Based Pharmacokinetic Model to Predict Disposition of CYP2D6 and CYP1A2 Metabolized Drugs in Pregnant Women. Drug Metabolism and Disposition 41, 801–813. doi:10.1124/dmd.112.050161

Keating, S. M., Waltemath, D., König, M., Zhang, F., Dräger, A., Chaouiya, C., et al. (2020). SBML Level 3: An extensible format for the exchange and reuse of biological models. Molecular systems biology 16, e9110. doi:10.15252/msb.20199110

Kerry, N., Somogyi, A., Bochner, F., and Mikus, G. (1994). The role of CYP2D6 in primary and secondary oxidative metabolism of dextromethorphan: In vitro studies using human liver microsomes. British Journal of Clinical Pharmacology 38, 243–248. doi:10.1111/j.1365-2125.1994.tb04348.x

Kibaly, C., Alderete, J. A., Liu, S. H., Nasef, H. S., Law, P.-Y., Evans, C. J., et al. (2021). Oxycodone in the Opioid Epidemic: High ‘Liking’, ‘Wanting’, and Abuse Liability. Cellular and Molecular Neurobiology 41, 899–926. doi:10.1007/s10571-020-01013-y

Kim, E.-Y., Shin, S.-G., and Shin, J.-G. (2017). Prediction and visualization of CYP2D6 genotype-based phenotype using clustering algorithms. Translational and Clinical Pharmacology 25, 147–152. doi:10.12793/tcp.2017.25.3.147

Köhler, D., Härtter, S., Fuchs, K., Sieghart, W., and Hiemke, C. (1997). CYP2D6 genotype and phenotyping by determination of dextromethorphan and metabolites in serum of healthy controls and of patients under psychotropic medication. Pharmacogenetics 7, 453–461. doi:10.1097/00008571-199712000-00003

[Dataset] König, M. (2021a). Matthiaskoenig/sbmlsim: 0.1.14 - SBML simulation made easy. https://zenodo.org/record/5531088. doi:10.5281/zenodo.5531088. Accessed: 2022-08-23

[Dataset] König, M. (2021b). Sbmlutils: Python utilities for SBML. https://zenodo.org/record/6599299. doi:10.5281/zenodo.5546603. Accessed: 2022-08-23

[Dataset] König, M. and Rodriguez, N. (2019). Matthiaskoenig/cy3sbml: Cy3sbml-v0.3.0 - SBML for Cytoscape. https://zenodo.org/record/3451319. doi:10.5281/zenodo.3451319. Accessed: 2022-08-23

König, M., Dräger, A., and Holzhütter, H.-G. (2012). CySBML: A Cytoscape plugin for SBML. Bioinformatics (Oxford, England) 28, 2402–2403. doi:10.1093/bioinformatics/bts432

Lenuzza, N., Duval, X., Nicolas, G., Thévenot, E., Job, S., Videau, O., et al. (2016). Safety and pharmacokinetics of the CIME combination of drugs and their metabolites after a single oral dosing in healthy volunteers. European journal of drug metabolism and pharmacokinetics 41, 125–138. doi:10.1007/s13318-014-0239-0

López, M., Guerrero, J., Jung–Cook, H., and Alonso, M. E. (2005). CYP2D6 genotype and phenotype determination in a Mexican Mestizo population. European Journal of Clinical Pharmacology 61, 749–754. doi:10.1007/s00228-005-0038-2

Lutz, J. D. and Isoherranen, N. (2012). Prediction of Relative In Vivo Metabolite Exposure from In Vitro Data Using Two Model Drugs: Dextromethorphan and Omeprazole. Drug Metabolism and Disposition 40, 159–168. doi:10.1124/dmd.111.042200

Machavaram, K. K., Endo-Tsukude, C., Terao, K., Gill, K. L., Hatley, O. J., Gardner, I., et al. (2019). Simulating the Impact of Elevated Levels of Interleukin-6 on the Pharmacokinetics of Various CYP450 Substrates in Patients with Neuromyelitis Optica or Neuromyelitis Optica Spectrum Disorders in Different Ethnic Populations. The AAPS journal 21, 42. doi:10.1208/s12248-019-0309-y

Mahgoub, A., Idle, J. R., Dring, L. G., Lancaster, R., and Smith, R. L. (1977). Polymorphic hydroxylation of Debrisoquine in man. Lancet (London, England) 2, 584–586. doi:10.1016/s0140-6736(77)91430-1

McGinnity, D. F., Parker, A. J., Soars, M., and Riley, R. J. (2000). Automated definition of the enzymology of drug oxidation by the major human drug metabolizing cytochrome P450s. Drug metabolism and disposition: the biological fate of chemicals 28, 1327–1334

Moghadamnia, A. A., Rostami-Hodjegan, A., Abdul-Manap, R., Wright, C. E., Morice, A. H., and Tucker, G. T. (2003). Physiologically based modelling of inhibition of metabolism and assessment of the relative potency of drug and metabolite: Dextromethorphan vs. dextrorphan using quinidine inhibition: Dextromethorphan vs. dextrorphan using quinidine inhibition. British Journal of Clinical Pharmacology 56, 57–67. doi:10.1046/j.1365-2125.2003.01853.x

Montané Jaime, L. K., Lalla, A., Steimer, W., and Gaedigk, A. (2013). Characterization of the CYP2D6 gene locus and metabolic activity in Indo-and Afro-Trinidadians: Discovery of novel allelic variants. Pharmacogenomics 14, 261–276. doi:10.2217/pgs.12.207

Myrand, S., Sekiguchi, K., Man, M., Lin, X., Tzeng, R.-Y., Teng, C.-H., et al. (2008). Pharmacokinetics/Genotype Associations for Major Cytochrome P450 Enzymes in Native and First-and Third-generation Japanese Populations: Comparison With Korean, Chinese, and Caucasian Populations. Clinical Pharmacology & Therapeutics 84, 347–361. doi:10.1038/sj.clpt.6100482

Nagai, N., Kawakubo, T., Kaneko, F., Ishii, M., Shinohara, R., Saito, Y., et al. (1996). Pharmacokinetics and polymorphic oxidation of dextromethorphan in a japanese population. Biopharmaceutics & Drug Disposition 17, 421–433. doi:10.1002/(SICI)1099-081X(199607)17:5〈421::AID-BDD421〉3.0.CO;2-9

Nakashima, D., Takama, H., Ogasawara, Y., Kawakami, T., Nishitoba, T., Hoshi, S., et al. (2007). Effect of Cinacalcet Hydrochloride, a New Calcimimetic Agent, on the Pharmacokinetics of Dextromethorphan: In Vitro and Clinical Studies. The Journal of Clinical Pharmacology 47, 1311–1319. doi:10.1177/0091270007304103

Ning, M., Duarte, J. D., Stevison, F., Isoherranen, N., Rubin, L. H., and Jeong, H. (2019). Determinants of Cytochrome P450 2D6 mRNA Levels in Healthy Human Liver Tissue. Clinical and Translational Science 12, 416–423. doi:10.1111/cts.12632

Nofziger, C. and Paulmichl, M. (2018). Accurately genotyping CYP2D6: Not for the faint of heart. Pharmacogenomics 19, 999–1002. doi:10.2217/pgs-2018-0105

Nyunt, M. M., Becker, S., MacFarland, R. T., Chee, P., Scarborough, R., Everts, S., et al. (2008). Pharmacokinetic Effect of AMD070, an Oral CXCR4 Antagonist, on CYP3A4 and CYP2D6 Substrates Midazolam and Dextromethorphan in Healthy Volunteers. JAIDS Journal of Acquired Immune Deficiency Syndromes 47, 559–565. doi:10.1097/QAI.0b013e3181627566

Oh, K.-S., Park, S.-J., Shinde, D. D., Shin, J.-G., and Kim, D.-H. (2012). High-sensitivity liquid chromatography-tandem mass spectrometry for the simultaneous determination of five drugs and their cytochrome P450-specific probe metabolites in human plasma. Journal of chromatography. B, Analytical technologies in the biomedical and life sciences 895–896, 56–64. doi:10.1016/j.jchromb.2012.03.014

Pope, L. E., Khalil, M. H., Berg, J. E., Stiles, M., Yakatan, G. J., and Sellers, E. M. (2004). Pharmacokinetics of Dextromethorphan After Single or Multiple Dosing in Combination With Quinidine in Extensive and Poor Metabolizers. The Journal of Clinical Pharmacology 44, 1132–1142. doi:10.1177/0091270004269521

Preskorn, S. H., Kane, C. P., Lobello, K., Nichols, A. I., Fayyad, R., Buckley, G., et al. (2013). Cytochrome P450 2D6 phenoconversion is common in patients being treated for depression: Implications for personalized medicine. The Journal of clinical psychiatry 74, 614–621. doi:10.4088/JCP.12m07807

Qiu, F., Liu, S., Miao, P., Zeng, J., Zhu, L., Zhao, T., et al. (2016). Effects of the Chinese herbal formula “Zuojin Pill” on the pharmacokinetics of dextromethorphan in healthy Chinese volunteers with CYP2D6*10 genotype. European journal of clinical pharmacology 72, 689–695. doi:10.1007/s00228-016-2048-7

Rau, T., Wohlleben, G., Wuttke, H., Thuerauf, N., Lunkenheimer, J., Lanczik, M., et al. (2004). CYP2D6 genotype: Impact on adverse effects and nonresponse during treatment with antidepressants-a pilot study. Clinical pharmacology and therapeutics 75, 386–393. doi:10.1016/j.clpt.2003.12.015

[Dataset] RNAO (2022). Nursing best practice guideline by nurses’ association of ontario: Appendix g: Tools for assessing anxiety, depression, and stress. https://bpgmobile.rnao.ca/sites/default/files/Appendix%20C%20Vein%20Anatomy.pdf. Accessed: 2022-08-23

Rougée, L. R. A., Mohutsky, M. A., Bedwell, D. W., Ruterbories, K. J., and Hall, S. D. (2016). The Impact of the Hepatocyte-to-Plasma pH Gradient on the Prediction of Hepatic Clearance and Drug-Drug Interactions for CYP2D6 Substrates. Drug metabolism and disposition: the biological fate of chemicals 44, 1819–1827. doi:10.1124/dmd.116.071761

Rüdesheim, S., Selzer, D., Fuhr, U., Schwab, M., and Lehr, T. (2022). Physiologically-based pharmacokinetic modeling of dextromethorphan to investigate interindividual variability within CYP2D6 activity score groups. CPT: Pharmacometrics & Systems Pharmacology n/a. doi:10.1002/psp4.12776

Sager, J. E., Lutz, J. D., Foti, R. S., Davis, C., Kunze, K. L., and Isoherranen, N. (2014). Fluoxetine-and norfluoxetine-mediated complex drug-drug interactions: In vitro to in vivo correlation of effects on CYP2D6, CYP2C19, and CYP3A4. Clinical pharmacology and therapeutics 95, 653–662. doi:10.1038/clpt.2014.50

Saravanakumar, A., Sadighi, A., Ryu, R., and Akhlaghi, F. (2019). Physicochemical Properties, Biotransformation, and Transport Pathways of Established and Newly Approved Medications: A Systematic Review of the Top 200 Most Prescribed Drugs vs. the FDA-Approved Drugs Between 2005 and 2016. Clinical Pharmacokinetics 58, 1281–1294. doi:10.1007/s40262-019-00750-8

Schadel, M., Wu, D., Otton, S. V., Kalow, W., and Sellers, E. M. (1995). Pharmacokinetics of dextromethorphan and metabolites in humans: Influence of the CYP2D6 phenotype and quinidine inhibition. Journal of Clinical Psychopharmacology 15, 263–269. doi:10.1097/00004714-199508000-00005

Schoedel, K. A., Pope, L. E., and Sellers, E. M. (2012). Randomized Open-Label Drug-Drug Interaction Trial of Dextromethorphan/Quinidine and Paroxetine in Healthy Volunteers:. Clinical Drug Investigation 32, 157–169. doi:10.2165/11599870-000000000-00000

Shah, R. R. and Smith, R. L. (2015). Addressing phenoconversion: The Achilles’ heel of personalized medicine: Impact of phenoconversion. British Journal of Clinical Pharmacology 79, 222–240. doi:10.1111/bcp.12441

Silva, A. R. and Dinis-Oliveira, R. J. (2020). Pharmacokinetics and pharmacodynamics of dextromethorphan: Clinical and forensic aspects. Drug Metabolism Reviews 52, 258–282. doi:10.1080/03602532.2020.1758712

Smith, L. P., Hucka, M., Hoops, S., Finney, A., Ginkel, M., Myers, C. J., et al. (2015). SBML Level 3 package: Hierarchical Model Composition, Version 1 Release 3. J Integr Bioinform 12, 268. doi:10.2390/biecoll-jib-2015-268

Somogyi, E. T., Bouteiller, J.-M., Glazier, J. A., König, M., Medley, J. K., Swat, M. H., et al. (2015). libRoadRunner: A high performance SBML simulation and analysis library. Bioinformatics (Oxford, England) 31, 3315–3321. doi:10.1093/bioinformatics/btv363

Steinberg, G. K., Bell, T. E., and Yenari, M. A. (1996). Dose escalation safety and tolerance study of the N-methyl-d-aspartate antagonist dextromethorphan in neurosurgery patients. Journal of Neurosurgery 84, 860–866. doi:10.3171/jns.1996.84.5.0860

Storelli, F., Desmeules, J., and Daali, Y. (2019a). Genotype-sensitive reversible and time-dependent CYP2D6 inhibition in human liver microsomes. Basic & clinical pharmacology & toxicology 124, 170–180. doi:10.1111/bcpt.13124

Storelli, F., Desmeules, J., and Daali, Y. (2019b). Physiologically-Based Pharmacokinetic Modeling for the Prediction of CYP2D6-Mediated Gene–Drug–Drug Interactions. CPT: Pharmacometrics & Systems Pharmacology 8, 567–576. doi:10.1002/psp4.12411

Strauch, K., Lutz, U., Bittner, N., and Lutz, W. K. (2009). Dose–response relationship for the pharmacokinetic interaction of grapefruit juice with dextromethorphan investigated by human urinary metabolite profiles. Food and Chemical Toxicology 47, 1928–1935. doi:10.1016/j.fct.2009.05.004

Takashima, T., Murase, S., Iwasaki, K., and Shimada, K. (2005). Evaluation of Dextromethorphan Metabolism Using Hepatocytes from CYP2D6 Poor and Extensive Metabolizers. Drug Metabolism and Pharmacokinetics 20, 177–182. doi:10.2133/dmpk.20.177

Tamminga, W., Wemer, J., Oosterhuis, B., de Zeeuw, R., de Leij, L., and Jonkman, J. (2001). The prevalence of CYP2D6 and CYP2C19 genotypes in a population of healthy Dutch volunteers. European Journal of Clinical Pharmacology 57, 717–722. doi:10.1007/s002280100359

Taylor, C. P., Traynelis, S. F., Siffert, J., Pope, L. E., and Matsumoto, R. R. (2016). Pharmacology of dextromethorphan: Relevance to dextromethorphan/quinidine (Nuedexta®) clinical use. Pharmacology & Therapeutics 164, 170–182. doi:10.1016/j.pharmthera.2016.04.010

Tennezé, L., Verstuyft, C., Becquemont, L., Poirier, J. M., Wilkinson, G. R., and Funck-Brentano, C. (1999). Assessment of CYP2D6 and CYP2C19 activity in vivo in humans: A cocktail study with dextromethorphan and chloroguanide alone and in combination. Clinical Pharmacology and Therapeutics 66, 582–588. doi:10.1053/cp.1999.v66.103401001

Tricco, A. C., Lillie, E., Zarin, W., O’Brien, K. K., Colquhoun, H., Levac, D., et al. (2018). PRISMA Extension for Scoping Reviews (PRISMA-ScR): Checklist and Explanation. Annals of Internal Medicine 169, 467–473. doi:10/gfd8vk

Vander, J. S., A. (2001). Human physiology: The mechanisms of body function. McGraw-Hill higher education

von Moltke, L. L., Greenblatt, D. J., Grassi, J. M., Granda, B. W., Venkatakrishnan, K., Schmider, J., et al. (1998). Multiple human cytochromes contribute to biotransformation of dextromethorphan in-vitro: Role of CYP2C9, CYP2C19, CYP2D6, and CYP3A. The Journal of Pharmacy and Pharmacology 50, 997–1004. doi:10.1111/j.2042-7158.1998.tb06914.x

[Dataset] Welsh, C., Xu, J., Smith, L., König, M., Choi, K., and Sauro, H. M. (2022). libRoadRunner 2.0: A High-Performance SBML Simulation and Analysis Library. doi:10.48550/arXiv.2203.01175

Whirl-Carrillo, M., Huddart, R., Gong, L., Sangkuhl, K., Thorn, C. F., Whaley, R., et al. (2021). An evidence-based framework for evaluating pharmacogenomics knowledge for personalized medicine. Clinical pharmacology and therapeutics 110, 563–572. doi:10.1002/cpt.2350

Wyen, C., Fuhr, U., Frank, D., Aarnoutse, R. E., Klaassen, T., Lazar, A., et al. (2008). Effect of an antiretroviral regimen containing ritonavir boosted lopinavir on intestinal and hepatic CYP3A, CYP2D6 and P-glycoprotein in HIV-infected patients. Clinical pharmacology and therapeutics 84, 75–82. doi:10.1038/sj.clpt.6100452

Yamazaki, T., Desai, A., Goldwater, R., Han, D., Howieson, C., Akhtar, S., et al. (2017). Pharmacokinetic Effects of Isavuconazole Coadministration With the Cytochrome P450 Enzyme Substrates Bupropion, Repaglinide, Caffeine, Dextromethorphan, and Methadone in Healthy Subjects. Clinical pharmacology in drug development 6, 54–65. doi:10.1002/cpdd.281

Yang, J., He, M. M., Niu, W., Wrighton, S. A., Li, L., Liu, Y., et al. (2012). Metabolic capabilities of cytochrome P450 enzymes in Chinese liver microsomes compared with those in Caucasian liver microsomes: Hepatic cytochromes P450 of Chinese and Caucasians. British Journal of Clinical Pharmacology 73, 268–284. doi:10.1111/j.1365-2125.2011.04076.x

Yu, A. and Haining, R. L. (2001). Comparative contribution to dextromethorphan metabolism by cytochrome P450 isoforms in vitro: Can dextromethorphan be used as a dual probe for both CTP2D6 and CYP3A activities? Drug Metabolism and Disposition: The Biological Fate of Chemicals 29, 1514–1520

Zackrisson, A. L., Lindblom, B., and Ahlner, J. (2010). High Frequency of Occurrence of CYP2D6 Gene Duplication/Multiduplication Indicating Ultrarapid Metabolism Among Suicide Cases. Clinical Pharmacology & Therapeutics 88, 354–359. doi:10.1038/clpt.2009.216

Zanger, U. M., Raimundo, S., and Eichelbaum, M. (2004). Cytochrome P450 2D6: Overview and update on pharmacology, genetics, biochemistry. Naunyn-Schmiedeberg’s archives of pharmacology 369, 23–37. doi:10.1007/s00210-003-0832-2

Zawertailo, L. A., Tyndale, R. F., Busto, U., and Sellers, E. M. (2010). Effect of metabolic blockade on the psychoactive effects of dextromethorphan. Human Psychopharmacology: Clinical and Experimental 25, 71–79. doi:10.1002/hup.1086

