## Supplementary Material for "Physiologically based pharmacokinetic (PBPK) modeling of the role of CYP2D6 polymorphism for metabolic phenotyping with dextromethorphan"

### 1 SUPPLEMENTARY TABLES AND FIGURES

| Allele | Activity score (AS) |
|---|---|
| *1 | 1.0 |
| *1x2 | 2.0 |
| *2 | 1.0 |
| *2x2 | 2.0 |
| *3 | 0 |
| *4 | 0 |
| *4x2 | 0 |
| *5 | 0 |
| *6 | 0 |
| *10 | 0.25 |
| *17 | 0.5 |
| *29 | 0.5 |
| *41 | 0.5 |

**Table S1. CYP2D6 allele-phenotype association**. The phenotype of CYP2D6 allele variants are characterized by the activity value adopted from PharmGKB.

| Activity Score | Sub-Saharan African | African American & Afro-Caribbean | European | Near Eastern | East Asian | Central & South Asian | American | Latino | Oceanian |
|---|---|---|---|---|---|---|---|---|---|
| 0 | 1.53 | 2.33 | 6.47 | 2.20 | 0.86 | 2.34 | 2.18 | 3.12 | 0.38 |
| 0.25 | 1.38 | 1.17 | 0.80 | 2.01 | 8.10 | 2.65 | 0.42 | 0.93 | 0.31 |
| 0.5 | 10.90 | 9.29 | 6.72 | 6.34 | 19.82 | 4.71 | 1.03 | 3.84 | 0.17 |
| 0.75 | 4.77 | 2.29 | 0.41 | 2.69 | 3.94 | 2.24 | 0.10 | 0.56 | 0.04 |
| 1 | 26.45 | 23.46 | 31.01 | 18.82 | 7.28 | 19.94 | 22.04 | 23.76 | 9.57 |
| 1.25 | 3.64 | 3.62 | 1.81 | 6.80 | 33.15 | 10.35 | 2.14 | 3.37 | 3.90 |
| 1.5 | 28.04 | 28.43 | 15.16 | 19.93 | 3.45 | 15.44 | 5.10 | 13.67 | 1.32 |
| 2 | 11.40 | 23.38 | 34.03 | 27.14 | 14.62 | 36.09 | 56.27 | 42.02 | 61.14 |
| 2.25 | 0.32 | 0.22 | 0.05 | 0.88 | 0.69 | 0.26 | 0.10 | 0.15 | 0.60 |
| 2.5 | 2.44 | 1.69 | 0.46 | 2.56 | 0.07 | 0.39 | 0.24 | 0.59 | 0.20 |
| 3 | 1.86 | 2.68 | 1.99 | 6.49 | 0.60 | 1.81 | 5.15 | 3.57 | 18.37 |
| 4 | 0.08 | 0.08 | 0.03 | 0.42 | 0.01 | 0.02 | 0.12 | 0.08 | 1.41 |
| **Phenotype** | | | | | | | | | |
| UM | 0.4 | 0.4 | 0.5 | 0.7 | 0.3 | 0.5 | 0.7 | 0.7 | 1.1 |
| EM | 73.7 | 76.6 | 73.8 | 78.2 | 66.3 | 77.5 | 83.3 | 79.1 | 88.5 |
| IM | 23.7 | 20.2 | 19.1 | 18.5 | 30.9 | 19.0 | 13.5 | 16.8 | 9.9 |
| PM | 2.2 | 2.7 | 6.7 | 2.7 | 2.6 | 2.9 | 2.4 | 3.5 | 0.5 |

**Table S2. Proportion [%] of CYP2D6 activity scores and simulated metabolic phenotypes by biogeographical group**.

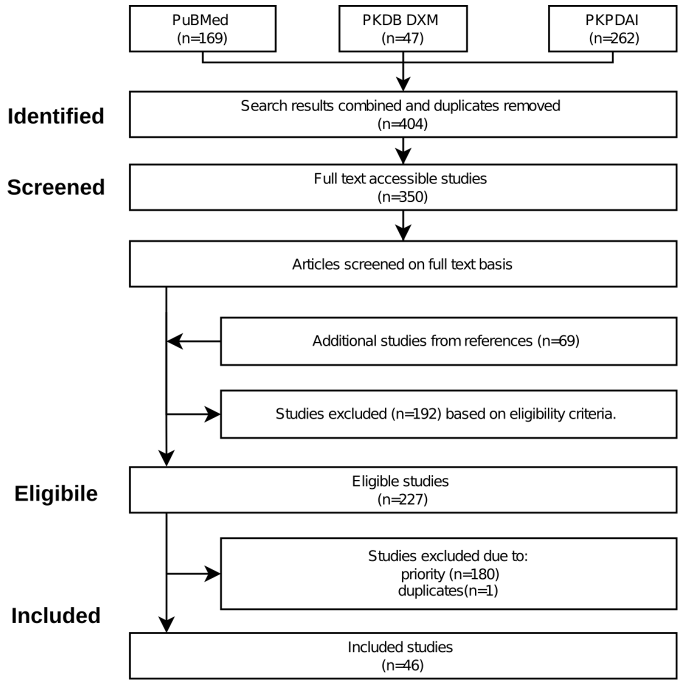


**Figure S1. PRISMA flow diagram.** Overview of the data selection for the pharmacokinetics dataset used in this work. PuBMed, PKDB, and PKPDAI were utilized for the literature search on DXM pharmacokinetics. Applied eligibility criteria resulted in 227 studies of which 46 were curated for this work. The process is described in detail in the Materials and Methods section.
